# A SUN1-ApoD feedback loop promotes cellular aging via microtubule-nuclear mechanotransduction

**DOI:** 10.64898/2026.09.08.750279

**Authors:** Mengqi Chen, Paige C. Wilson, Wenyao Yang, Yue Ma, Yutao Li, Xinluan Wang, Richard L. Bennett, Jonathan D. Licht, Gregg G. Gundersen, Wakam Chang

**Affiliations:** MOE Frontier Science Centre for Precision Oncology, University of Macau; Taipa, Macau, China; Department of Biomedical Sciences, Faculty of Medicine, University of Macau; Taipa, Macau, China; Guangdong-Hong Kong-Macao Joint Laboratory for New Drug Screening, University of Macau; Taipa, Macau, China; Department of Biological Sciences, Columbia University; New York, NY, USA; Translational Medicine R&D Center, Institute of Biomedical and Health Engineering, Shenzhen Institutes of Advanced Technology, Chinese Academy of Sciences; Shenzhen, 518000, China; Musculoskeletal Research Laboratory, Department of Orthopaedics and Traumatology, The Chinese University of Hong Kong; Kowloon, Hong Kong, China; Department of Epigenetics, Van Andel Institute, Grand Rapids, MI 49503, USA; Department of Pathology and Cell Biology, Vagelos College of Physicians and Surgeons, Columbia University; New York, NY, USA

**Keywords:** nesprin-2, LINC complex, nuclear envelope, cell polarity, gene expression

## Abstract

Alterations in nuclear envelope proteins such as SUN1 and prelamin A are known hallmarks of cellular aging, but how they cause aging phenotypes beyond the expressing cells is unknown. With a tissue culture version of parabiosis, we found that aged fibroblasts secrete an activity that induces aging-related polarity defects in young fibroblasts. Secretomics identified the factor as apolipoprotein D (ApoD), a circulating protein whose levels are known to rise with age. ApoD increased SUN1 levels and disrupted polarity in fibroblasts via SUN1-promoted coupling of microtubules to the nucleus. In turn, elevated SUN1 enhanced ApoD expression and secretion, forming a feedback loop that promotes acquisition of aged phenotypes. In mice, ApoD induced aging-related phenotypes in muscle. Elevated ApoD expression required coupling between SUN1, its outer nuclear membrane binding partner nesprin-2, and microtubules. Broader analysis revealed that this pathway regulates hundreds of genes. These findings define microtubule-nuclear coupling as a direct mechanotransduction pathway controlling gene expression and promoting aging-associated phenotypes.

## Main Text

Aging is characterized by cellular hallmarks that are shared across the phylogenetic landscape^1^. A number of these include processes related to the nucleus, such as genomic instability, telomere attrition, epigenetic changes, and cellular senescence. Recent progress suggests that alterations in the nucleus itself may be a new hallmark of aging^2,3^. Direct evidence comes from premature aging (progeroid) syndromes such as Hutchison-Gilford progeria syndrome (HGPS) and mandibuloacral dysplasia B (MDB), in which the accumulation of farnesylated prelamin A alters the mechanical properties of the nucleus^4,5^.

While prelamin A does not characteristically accumulate in physiologically aged cells^6,7^, they exhibit many of the same nuclear changes found in progeroid syndromes^8,9^. In fact, there is strong evidence linking both premature and physiological aging to elevated levels of the inner nuclear membrane protein SUN1^10–16^. SUN1 binds to nesprin/KASH proteins in the outer nuclear membrane to form the linker of nucleoskeleton and cytoskeleton (LINC) complex^17,18^. This complex transmits cytoskeleton forces to move the nucleus^19–22^, regulate nuclear pores^23^, alter the epigenome^24,25^, activate gene expression^26–28^, and facilitate DNA repair^29,30^. Elevated SUN1 is critical to premature aging phenotypes, as crossing mouse models of premature aging with *Sun1* knockout mice extends their lifespan with physiological improvement in heart, bone, and other tissues^13^. Studies of cultured fibroblasts from individuals with premature aging syndromes or from aged individuals show that elevated SUN1 levels enhance interaction of the nucleus with microtubules (MTs) and reduce its interaction with actin filaments^10,16^. This switch in cytoskeletal engagement prevents actin-dependent movement of the nucleus and centrosome orientation important for generating front-rear polarity during migration. Artificially elevating SUN1 levels in young human fibroblasts (HFs) or immortalized mouse NIH3T3 fibroblasts is sufficient to promote MT-nuclear interaction and inhibit actin-nuclear coupling^10,16^.

The mechanistic basis for elevated SUN1 levels and their contribution to aging phenotypes across numerous tissues is unknown. Here, we use the inability of aged fibroblasts to generate actin-dependent nuclear movement and centrosome orientation (“cell polarization”) as assays to test whether SUN1 levels are affected by secreted factors using a modified parabiosis approach. We identify ApoD as a factor selectively secreted by aged fibroblasts that inhibits polarization of young fibroblasts and elevates SUN1 levels. SUN1 in turn promotes ApoD expression and secretion. Strikingly, the regulation of the secreted factors by SUN1 occurs through a pathway of direct mechanical regulation of gene expression by MTs rather than previously described pathways involving actin contractility^31,32^.

### Aged cells secrete an activity that inhibits fibroblast polarity

Recent studies have shown that aging accelerates markedly between 50 and 60 years of age^33,34^. Fibroblasts isolated from individuals also exhibited an abrupt transition from a normal to an abnormal polarized state during this period^10^. This biphasic switch suggests an underlying synchronization mechanism, potentially mediated by factors that coordinate cellular responses through cell-cell communication. We therefore used a tissue culture version of parabiosis experiments, which have revealed soluble factors contributing to young or aged phenotypes in animals^35–39^. We treated aged HFs with conditioned medium (CM) from serum-starved young HFs and conversely, young HFs with CM from serum-starved aged HFs and assessed nuclear movement and centrosome orientation (Fig. 1A) in response to lysophosphatidic acid (LPA) stimulation, which activates these processes in fibroblasts^40,41^. CM from aged cells significantly impaired rearward nuclear positioning (Fig. 1B, C) and centrosome orientation (Fig. 1B, D) of young HFs. In contrast, neither CM from young or aged cells or fresh media improved rearward nuclear positioning (Fig. 1C) or centrosome orientation in aged HFs (Fig. 1D). Thus, aged HFs release an activity capable of inhibiting cell polarization in young HFs.

**Fig. 1.**
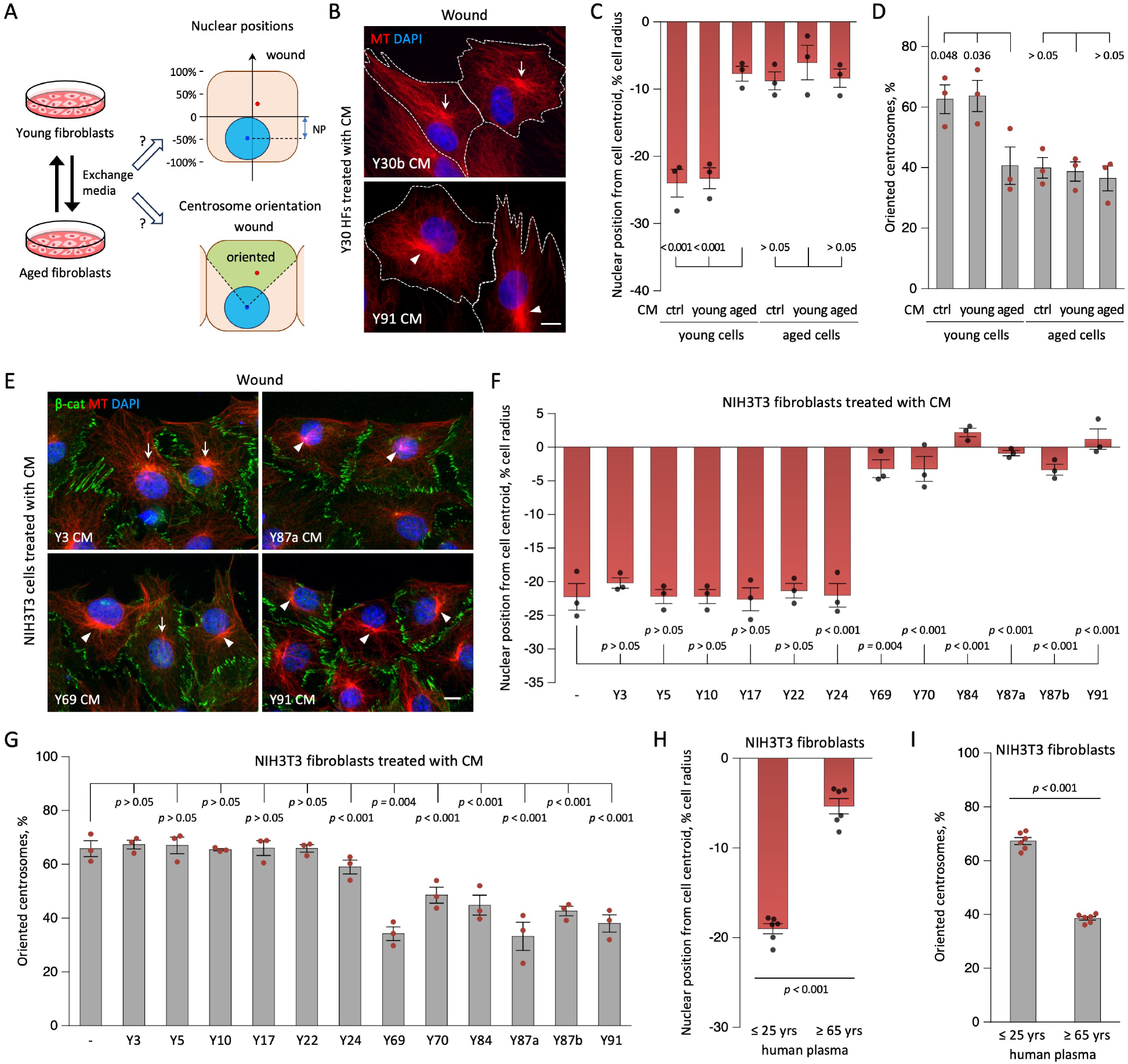
An inhibitor of cell polarity is released from aged HFs and is present in plasma from aged individuals. (**A**) Schematics depicting the experimental design (left) and how nuclear position (right, top) and centrosomal orientation (right bottom) were assessed. For nuclear position, the distance between the nuclear and cell centroids along the front-back axis was determined and normalized to cell radius (-values are rear of the cell centroid; + values are forward). For centrosome orientation, the percentage of cells with centrosomes in the green area was determined. (**B**) Representative images of LPA-stimulated HFs from young individuals treated with CM from HFs of young and aged donors and stained for MTs and nuclei (DAPI). Note the rearward positioned nuclei and oriented centrosome (between the nucleus and cell front, arrows) in the top panel and the central positioned nuclei and unoriented centrosomes (arrowheads) in the bottom panel. (**C, D**) Quantification of nuclear positions (C) and centrosome orientation (D) in HFs from young and aged individuals treated with fresh serum-free medium (control), CM from young, and CM from aged HFs. (**E**) Representative images of LPA-stimulated NIH3T3 fibroblasts treated with CMs from the indicated HFs (Y, age in years) and stained for MT, DAPI and β-catenin (to outline cell edges). Arrows, orientated centrosome; arrowheads unoriented centrosomes. (**F**,**G**) Quantification of nuclear position (F) and centrosome orientation (G) in LPA-stimulated for NIH3T3 fibroblasts treated as in E. (**H, I**) Quantification of nuclear position (H) and centrosome orientation (I) of LPA-stimulated NIH3T3 fibroblasts treated with plasma from six young and six aged donors. Each point corresponds to data from an individual donor. Bars, 10 µm. Values in C, D, F and G are mean ± SEM and points represent individual determinations from *N* = 3 experiments, *n* ≥ 90 cells; H, I: *N* = 6 donors, *n* ≥ 30 cells). *P* values were calculated with 1-way ANOVA with Tukey’s (C, D) or Dunnett’s post hoc test (F, G), or unpaired two-tailed *t*-test (H, I).

Immortalized mouse NIH3T3 fibroblasts provide a reproducible and well-studied system for studying the acquisition of cell polarity^41–46^ and are more amenable to manipulation than primary HFs. We found that CM from HFs from individuals ≥ 69 years (Table S1), previously shown to be defective in the establishment of cell polarity^10^, inhibited LPA-induced posterior nuclear positioning and centrosome orientation, whereas CM from young cells (3-23 years) had no effect (Fig. 1E-G). Thus, aged HFs release a factor that inhibits the acquisition of polarity in both human and mouse fibroblasts. Notably, cell polarity establishment was also suppressed by human plasma from individuals ≥ 65 years, but not those from individuals ≤ 25 years (Fig. 1H, I), underscoring the physiological relevance of the finding.

Fibroblasts from individuals with premature aging syndromes such as HGPS and MDB are also defective in repositioning their nucleus rearward and orienting their centrosomes^10,16^. We found that CM from HGPS fibroblasts, but not those from chronologically age-matched wild-type controls, inhibited the cell polarization parameters in NIH3T3 fibroblasts (Fig. S1A, B). The polarity inhibitory activity could also be induced in the CM from NIH3T3 fibroblasts expressing myc-progerin, which upregulates SUN1^10^, but not CMs of cells expressing empty vector or myc-lamin A (Fig. S1C, D). Thus, the polarity inhibiting factor is characteristic of both physiological and premature aged fibroblasts and is conserved between human and mice.

Senescent cells are known to release a variety of cytokines as part of the senescence-associated secretory phenotype (SASP)^47^. However, previously we did not detect a difference in the levels of two senescence markers, DNA damage and β-galactosidase activity, between the young and aged HFs used in this study^10^. Moreover, when we cultured young HFs until they became senescent, they polarized normally^10^. The CM from these senescent young HFs did not affect polarization of NIH3T3 fibroblasts (Fig. S2), suggesting that the observed polarity inhibitory factor is distinct from SASP and is specifically associated with chronological aging.

### ApoD is a polarity inhibitor released by aged cells

To identify the cell polarity inhibitor released by aged HFs, we first characterized its biochemical properties. We used the inhibition of rearward nuclear positioning in NIH3T3 fibroblasts as a readout for altered cell polarity and tested CM from at least two HFs in each case. The polarity inhibitory activity of CM from aged HFs was not dialyzable (12.5-kDa cutoff) and was abolished by heat inactivation (Fig. 2A, S3A, B), suggesting a macromolecular nature. The inhibitor remained soluble after ultracentrifugation and was unaffected by RNase digestion or charcoal adsorption (Fig. 2A, Fig. S3C-E) but was inactivated by trypsin and proteinase K (Fig. 2A, S3E). These results indicate that the inhibitor was a protein and excluded involvement of exosomes, RNA, or lipids.

**Fig. 2.**
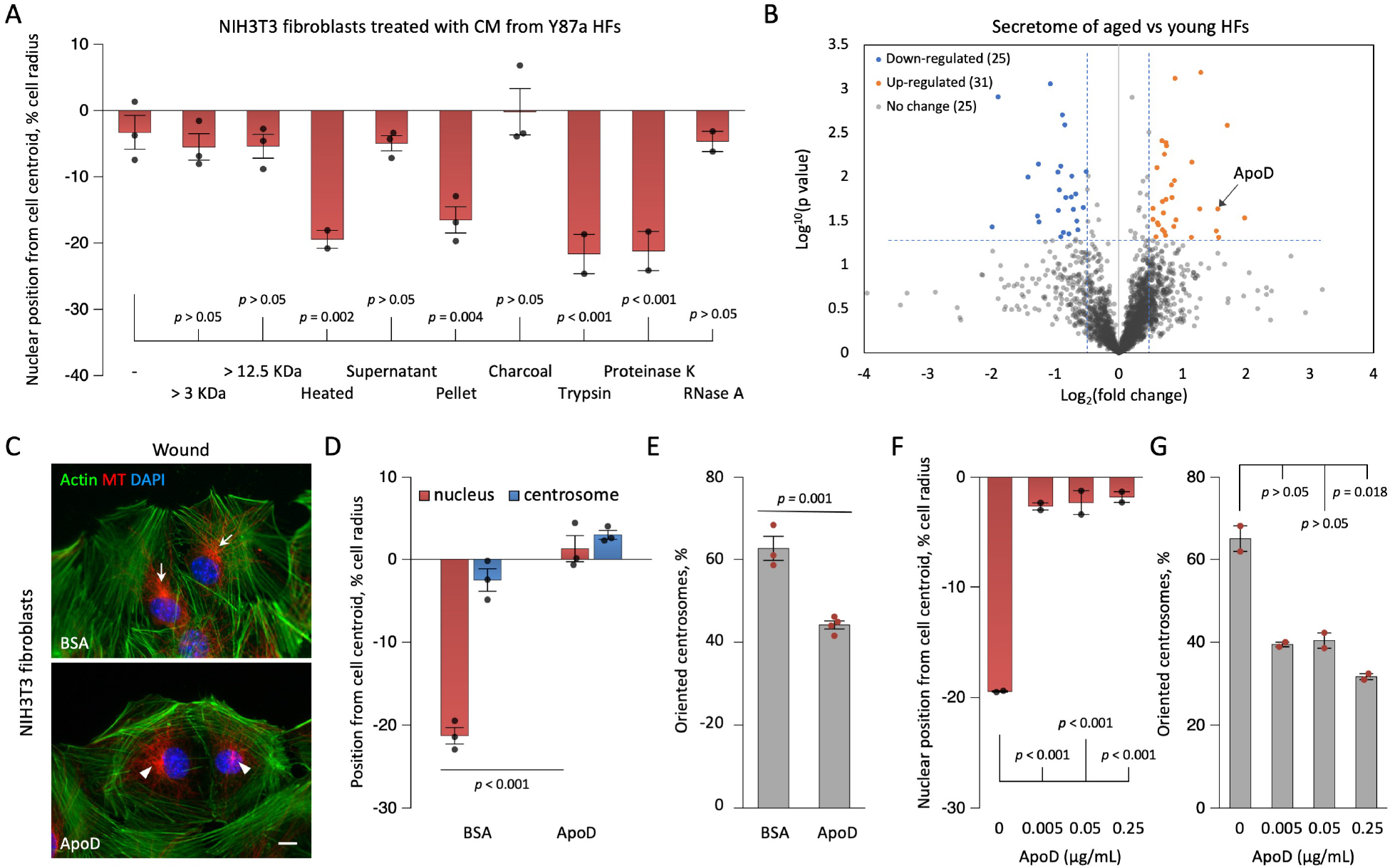
ApoD is a polarity inhibitor in CM from aged HFs. (**A**) Nuclear position in LPA-stimulated NIH3T3 fibroblasts treated with CM from aged HFs after the indicated biochemical treatments: >3 KDa and >12.5 KDa, retentate after dialysis with a membrane with the indicated molecular weight cutoff; Heated, 100 °C, 10 min; Supernatant and Pellet, fractions after centrifugation 100,000 ×*g*, 30 min; Charcoal, treatment with charcoal; Trypsin and Proteinase K: protein digestion; RNases A: RNA digestion. Note the loss of inhibitory activity after heat inactivation and protease treatment. See Fig. S3 for additional details. (**B**) Volcano plot showing upregulated (orange) and downregulated (blue) proteins in CMs from aged vs. young HFs. ApoD is among the highest upregulated proteins in CM from aged fibroblasts. (**C**) Representative images of LPA-stimulated NIH3T3 fibroblast pretreated with ApoD (0.1 µg/mL, 24 h) or BSA as a control and stained for F-actin (phalloidin), MTs and nuclei (DAPI). Arrows, orientated centrosome; arrowheads, unoriented centrosomes. Bar in C, 10 µm. (**D, E**) Quantification of nuclear position (D) and centrosome orientation (E) for cells treated as in C. (**F, G**) Quantification of nuclear position (F) and centrosome orientation (F) for cells treated with ApoD at the indicated concentration. Values are mean ± SEM (A: *N* = 2-3 experiments, *n* ≥ 60 cells; D, E: *N* = 3 experiments, n ≥ 90 cells; F, G: *N* = 2 experiments, *n* ≥ 60 cells). *P* values were calculated with 1-way ANOVA with Dunnett’s post hoc test (A, F, G) or unpaired two-tailed *t*-test (D-E).

The polarity inhibitory activity in CM from aged HFs was detectable after diluting it 20-25-fold (Fig. S4A, B). This result suggested it was present in a significant concentration that would be amenable to proteomic identification, particularly given that the CM lacked serum proteins. We used tandem mass tag mass spectrometry to compare proteins in CM from three young and three aged HFs. Of the 2,519 proteins identified (Source Data S1), 31 proteins were upregulated and 25 were downregulated in aged HFs compared to young HFs (criteria: *p* < 0.05 and log2 fold change ≥ 0.5) (Fig. 2B). Among the differentially detected proteins, we focused on ApoD, as it had one of the highest fold-increases (2.9-fold) in CM from aged cells (Fig. 2B), was previously reported to increase during aging^48^, and is also a normal constituent of blood/plasma^49^. We confirmed that ApoD levels were elevated in the plasma of aged individuals compared with those of young individuals (Fig. S4C). Treatment of NIH3T3 fibroblasts with recombinant ApoD inhibited rearward nuclear positioning (Fig. 2C, D) and centrosome orientation (Fig. 2C, E). Inhibition was observed over a range of ApoD concentrations (0.03–50 µg/mL) reported in plasma (Fig. 2E, F)^48,50^. Thus, ApoD is a polarity inhibitor selectively secreted by aged cells.

### ApoD inhibits cell polarity through SUN1 and MTs

To elucidate how ApoD inhibited cell polarization, we first tested the simple explanation that it might sequester LPA and block its activity. LPA activates Rho GTPase to assemble filamentous actin (F-actin) in starved fibroblasts, and this response to LPA was indistinguishable in cells treated with or without ApoD (Fig. S5). Short treatments (2 h) with ApoD did not inhibit LPA-induced rearward nuclear positioning (Fig. 3A) or centrosome orientation (Fig. 3B). Similarly. short-term exposure to CM from aged HFs did not impair cell polarity (Fig. S6A, B). Cells pretreated with ApoD for one day failed to polarize even after ApoD was removed for an additional day (Fig. 3C, D). We also added ApoD to the cells before serum starvation and found that its effect persisted for two days and it remained effective in the presence of serum (Fig. S6C, D). These results confirm that ApoD does not function by inhibiting the response to LPA and suggest a longer-term change in the cells is responsible.

**Fig. 3.**
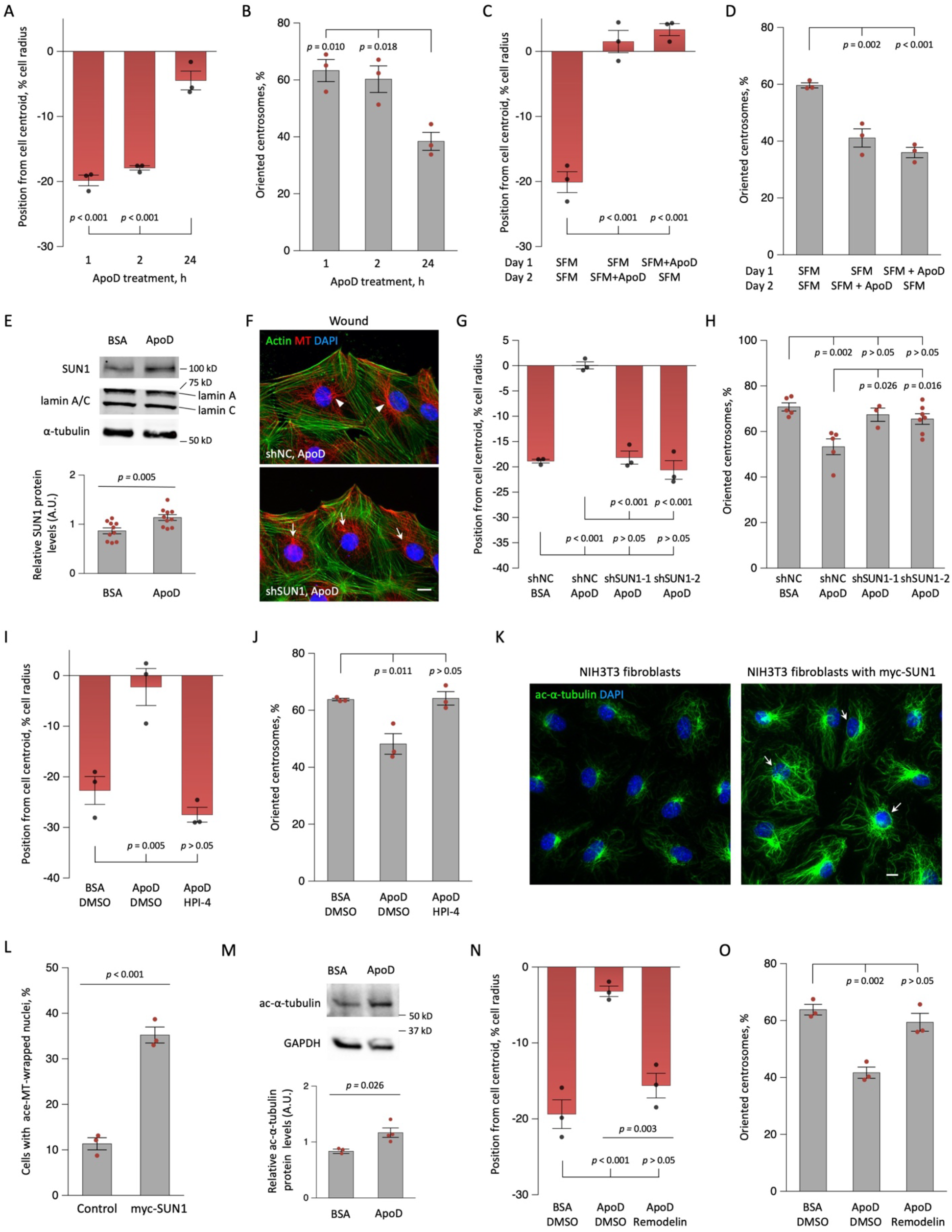
ApoD inhibits nuclear movement through SUN1 and MTs. (**A, B**) Nuclear position (A) and centrosome orientation (B) in LPA-stimulated NIH3T3 fibroblasts treated with ApoD (0.1 µg/mL) for the indicated periods. (**C, D)** Nuclear position (C) and centrosome orientation (D) in LPA-stimulated NIH3T3 fibroblasts treated with ApoD for 1 day and then washed out. SFM: serum-free medium. (**E**) Immunoblot (top) and quantification (bottom) showing upregulation of SUN1, but not progerin (migrates between lamin A and C bands), in lysates of NIH3T3 fibroblasts treated with ApoD (0.1 µg/mL 24 h). Tubulin is a loading control. (**F**) Representative images of LPA-stimulated NIH3T3 fibroblasts treated with ApoD and SUN1 or non-coding (NC) shRNAs and stained for actin (phalloidin), MT and nuclei (DAPI). Arrows, orientated centrosome; arrowheads, unoriented centrosomes. (**G, H**) Nuclear position (G) and centrosomal orientation (H) for cells treated as in F. (**I, J**) Nuclear position (I) and centrosome orientation (J) in LPA-stimulated NIH3T3 fibroblasts treated with ApoD or BSA (control) and the dynein inhibitor HPI-4 or DMSO vehicle. (**K**) Representative images of acetylated (ac) α-tubulin in NIH3T3 fibroblasts without and with expression of myc-SUN1. Arrows, nuclei wrapped by acetylated MT. (**L**) Quantification of nuclei wrapped by acetylated MT in K. (**M**) Immunoblot (top) and quantification (bottom) of acetylated α-tubulin in NIH3T3 fibroblasts treated with 0.1 µg/mL BSA or ApoD. (**N, O**) Nuclear position (N) and centrosome orientation (O) in LPA-stimulated NIH3T3 fibroblasts treated with BSA, ApoD, or ApoD together with 50 µM Remodelin. Values are mean ± SEM (E: *N* = 10 experiments; L: *N* = 3 experiments, *n* ≥ 300 cells; M: *N* ≥ 3 experiments; other panels: *N* ≥ 3 experiments, *n* ≥ 90 cells). *p* values were calculated with 1-way ANOVA with Tukey’s post hoc test (except E, L, M) or unpaired two-tailed *t*-test (E, L, M).

Since progerin expression and elevated SUN1 are linked to aging and interfere with nuclear movement like ApoD^10,13^, we determined whether ApoD affected their expression. ApoD treatment did not induce progerin but reproducibly increased SUN1 protein levels (Fig. 3E). While *Sun1* mRNA expression remained unchanged by ApoD (Fig. S7A), SUN1 protein stability was altered, as shown by a cycloheximide chase assay (Fig. S7B, C).

Given that elevated SUN1 interferes with SUN2-supported actin-dependent nuclear movement^10,20,51^, we asked whether the ApoD-stimulated increase in SUN1 accounted for the inhibition of cell polarity. Depletion of SUN1 using two shRNAs (Fig. S7D) prevented ApoD from inhibiting rearward nuclear positioning and centrosome orientation (Fig. 3F-H). SUN1 blocks actin-dependent nuclear movement by promoting nesprin-2G association with MTs rather than actin. This effect can be reversed by inhibiting the dynein motor protein that connects nesprin-2 to MTs^10,20,51^. Accordingly, treatment of cells with a dynein inhibitor, HPI-4, blocked the inhibitory effect of ApoD on polarity (Fig. 3I, J). SUN1 expression increases α-tubulin acetylation^51^ and these acetylated MTs accumulated around the nucleus (Fig. 3K, L). ApoD treatment also increased α-tubulin acetylation (Fig. 3M), and reducing acetylation using remodelin^51–53^ also blocked the ability of ApoD to inhibit cell polarity (Fig. 3N, O). These results indicate that ApoD inhibits actin-dependent nuclear movement by upregulating SUN1 protein level and promoting microtubule association with the nucleus.

### SUN1 expression induces ApoD secretion

ApoD levels increase with age and are elevated by stress^48,54,55^. We hypothesized that the elevated SUN1 levels associated with aging^10,13^ might stress the nucleus by changing the balance of nuclear connection to MTs *versus* actin, thereby affecting ApoD expression and/or secretion. We expressed myc-tagged wild-type SUN1 (WT) or a SUN1 C759A mutant in NIH3T3 fibroblasts (Fig. S7E) and assayed ApoD levels in CM. The SUN1 C759A mutant prevents covalent interaction with nesprins^56^ and cannot enhance microtubule-nuclear interactions^51^. ApoD protein (Fig. 4A) as well as mRNA (Fig. 4B) levels were substantially elevated by expression of SUN1 WT compared to SUN1 C759A or non-expressing controls. Moreover, reducing SUN1 levels by shRNA depletion reduced ApoD secretion in NIH3T3 fibroblasts (Fig. 4C). As expected, CM from SUN1 WT expressing cells, but not SUN1 C759A cells, inhibited nuclear positioning and centrosome orientation (Fig. 4D, E). These results indicate that ApoD expression and secretion are modulated by SUN1 levels and that this regulation likely reflects a microtubule-nuclear mechanotransductive pathway given that the SUN1 C759A mutant lacked this activity.

**Fig. 4.**
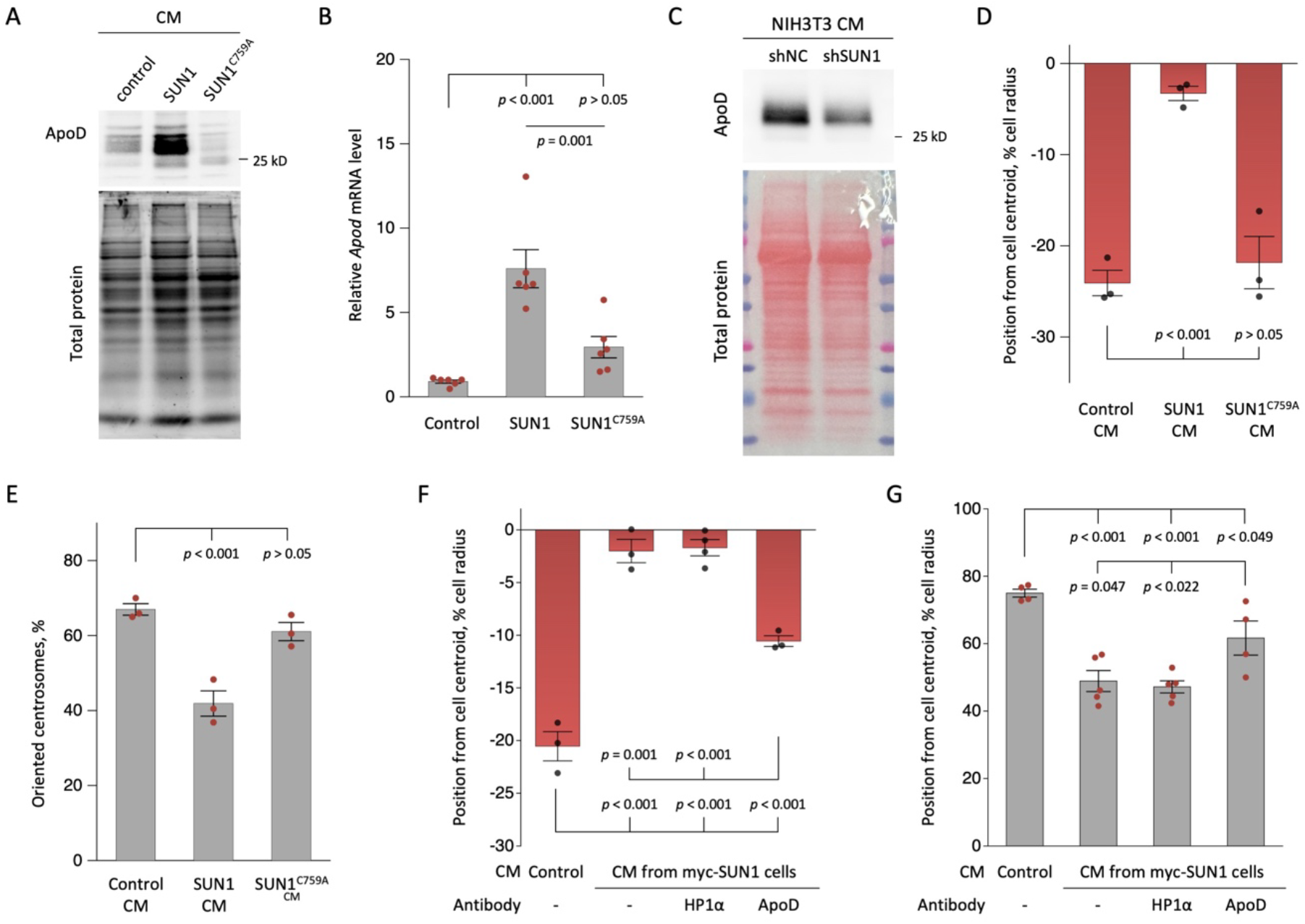
SUN1 regulates ApoD expression through its interaction with a nesprin. (**A**) Immunoblot of ApoD and Coomassie blue staining (total protein) in concentrated CMs from control and cells expressing myc-SUN1 and myc-SUN1^C759A^. (**B**) Quantification of *Apod* mRNA levels in cells expressing indicated protein. (**C**) Top: Immunoblot showing the levels of ApoD in concentrated CMs from NIH3T3 fibroblasts expressing the indicated shRNAs. Bottom: Ponceau S staining showing total protein. (**D, E**) Nuclear position (D) and centrosome orientation (E) in LPA-stimulated NIH3T3 fibroblasts treated with CM from cells expressing the indicated proteins. (**F, G**) Nuclear position (F) and centrosome orientation (G) of LPA-stimulated NIH3T3 fibroblasts treated with CM from cells expressing myc-SUN1 and ApoD-specific antibody or control HP-1α antibody. Values are mean ± SEM (B: *N* = 6 experiments; D-G: *N* ≥ 3 experiments, *n* ≥ 90 cells). *p* values were calculated with 1-way ANOVA with Tukey’s (B, F, G) or Dunnett’s (D, E) post hoc test.

We next asked if ApoD is the sole polarity inhibitor induced by SUN1. We attempted to immunodeplete ApoD from CM without success. Alternatively, we directly added an ApoD-specific antibody to CM from SUN1 expressing cells to neutralize ApoD. The ApoD antibody partially blocked the inhibitory effect of CM from SUN1 expressing cells, whereas an irrelevant antibody (HP1α) had no effect (Fig. 4F, G). This suggests that ApoD is the major polarity inhibitor secreted by SUN1-overexpressing cells. Together, these results suggest that a SUN1 LINC complex regulates ApoD expression and secretion and elicits a positive feedback loop between SUN1 protein levels and ApoD secretion.

### ApoD is one of a group of genes regulated by a direct MT-nuclear gene expression pathway

Given that SUN1 regulation of ApoD expression appeared to require a covalent connection to nesprins, and prior work showing that SUN1 promotes nesprin-2 interaction with MTs^10,20,51^, we next tested whether SUN1 regulated gene expression through a LINC complex connection to MT. We knocked down nesprin-2G (Fig. S8A), the major mechanoresponsive nesprin connecting actin filaments and MTs to the nucleus in NIH3T3 fibroblasts^10,20,51^, and performed transcriptomic analysis by RNA-Seq. Nesprin-2G knockdown altered the expression of 356 genes (criteria: *p* < 0.05, fold change ≥ 1.5): 276 were downregulated and 80 were upregulated (Fig. 5A, Source Data S2). Among the downregulated genes, *Apod* was one of the most affected with its level reduced 3.8-fold.

**Fig. 5.**
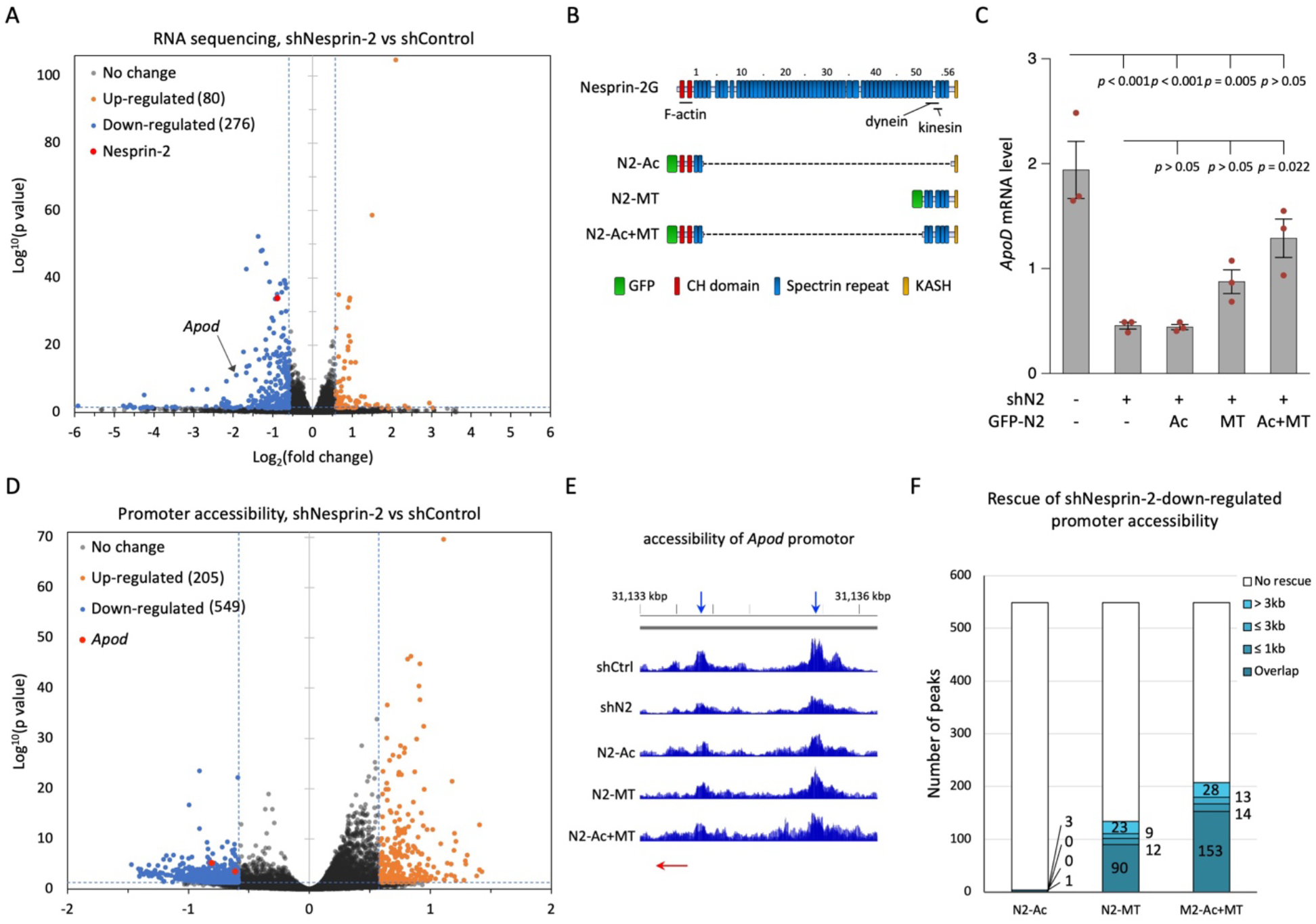
Nesprin-2 regulates gene expression through MTs. (**A**) Volcano plot for both upregulated (orange) and downregulated (blue) differentially regulated genes comparing NIH3T3 fibroblasts expressing control (shControl) and nesprin-2 (shNesprin-2) shRNAs. (**B**) Diagram depicting the domain structures of nesprin-2G and the fragments. Nesprin-2G binds actin through its N-terminal calponin homology (CH) domains and MTs through association with dynein and kinesin-1 motors. (**C**) *Apod* mRNA level in cells expressing shN2 and the indicated GFP-tagged nesprin-2 fragments. **(D)** Volcano plot from ATAC-seq analysis showing promoter regions with increased accessibility (orange, FDR < 0.05, fold change > 1.5) and decreased accessibility (blue) in NIH3T3 cells expressing shControl versus shNesprin-2. Two peaks in the *Apod* promoter region were identified. (**E**) Diagram showing the accessibility at the *Apod* promoter region from 31,133,000 to 31,136,500 bp on chromosome 16. Transcription start site is at 31,133,653. Blue arrows, peaks down regulated by shNesprin-2. Red arrow, transcription start site and direction. See Fig. S9A for the accessibility of the *Apod* gene body. **(F)** Number of downregulated promoter regions in H whose accessibility was increased by expression of the indicated nesprin-2 fragments. The regions were classified according to the distances between the shNeprin-2-downregulated peak and the peak induced by the indicated nesprin-2 fragment. “Overlap” indicates direct overlap between the corresponding peaks; ≤ 1 kb, ≤ 3 kb, and > 3 kb denote the distance between corresponding peaks; “No rescue” indicates regions where accessibility was not restored. Values in C are mean ± SEM (*N* = 3 biological repeats) and *p* values were calculated with 1-way ANOVA with Tukey’s post hoc test.

We next tested whether expression of *Apod* required nesprin-2G connection to actin filaments or MTs by performing RNA-Seq on NIH3T3 fibroblasts with nesprin-2G knockdown and re-expressed GFP-tagged fragments of nesprin-2G known to interact with actin filaments via calponin homology domains (N2-Ac), with MTs via MT-motors (N2-MT), or with both (N2-Ac+MT) (Fig. 5B). All fragments contained the KASH domain for nuclear targeting and SUN protein binding and were previously shown to functionally support actin- or MT-dependent nuclear movements in various contexts^19,20,31,57^. The re-expressed proteins exhibited predicted sizes (Fig. S8A) and localized to the nuclear envelope (Fig. S8B). The downregulated expression of *Apod* in the nesprin-2G depleted cells was elevated by re-expression of N2-MT (although not statistically significant) but was fully restored by re-expression of the N2-Ac+MT fragment (Fig. 5C). In the larger set of genes differentially regulated by nesprin-2G (Source Data S2), re-expression of N2-MT altered the expression of 195 genes (Fig. S8C). Most of the upregulated genes (81 of 150) were ones that downregulated by nesprin-2G knockdown, corresponding to a 29.3% rescue (Fig. S8D). Re-expression of the N2-Ac+MT fragment rescued almost half of the nesprin-2 responsive genes (Fig. S8D). In contrast, re-expression of N2-Ac had minimal effects on nesprin-2 responsive genes rescuing only 5 (1.8%) of them (Fig. S8D). Notably, the majority of N2-MT rescued genes (69 of 93, 74.2%) were also rescued by N2-Ac+MT (Fig. S8E), showing the robustness of the analysis and the critical requirement for MTs. Gene set enrichment analysis (GSEA) of biological processes (BP) among N2-MT-rescued genes revealed enrichment of pathways involved in host-pathogen interaction and cytokine signaling (Table S2). Taken together, our results suggest an unexpected role of nesprin-2 and MT mechanical coupling in regulating *Apod* expression and a broader set of genes.

We next conducted ATAC-Seq on the nesprin-2G depleted NIH3T3 fibroblasts and those re-expressing nesprin-2G fragments (Source Data S3). Nesprin-2 depletion resulted in an overall reduction in accessibility with 549 promoter-associated peaks less accessible and 205 more accessible (Fig. 5D). The promoter region of the *Apod* gene became less accessible in nesprin-2 depleted cells and its accessibility was restored by re-expressing either the N2-MT or N2-MT+Ac fragment, but not the N2-Ac fragment (Fig. 5E, S9A). For the broader set of nesprin-2 responsive peaks, the accessibility of 111 (24.4%) and 180 (37.9%) were restored

(using the emergence of a differential accessibility peak within 3 kb of the previously downregulated peak as the criterion) by N2-MT and N2-Ac+MT, respectively (Fig. 5F). In contrast, N2-Ac expression led to increased accessibility in only a single promoter (0.2%) (Fig. 5F). For regions whose accessibility was increased following nesprin-2 depletion, we observed a similar rescue pattern with the N-2 MT or N2-Ac+MT fragments (Fig. S9B). Together, these results support a model in which the attachment of MTs to the N2-SUN1 LINC complex regulates gene expression.

### Elevated SUN1 expression and ApoD treatment induce multiple markers of cellular aging

We surveyed other aging hallmarks to determine whether the altered MT nuclear mechanotransduction in cells with elevated SUN1 and ApoD stimulated aging phenotypes besides defective cell polarity. Expression of WT SUN1 in fibroblasts cultured for extended intervals (10-14 d) displayed reduced lamin B1 expression (Fig. 6A), a marker of aging^58^. Consistent with the need for force transmission through nesprin-2, SUN1 C759A did not cause lamin B1 reduction. Expression of SUN1 WT, but not SUN1 C759A, in NIH3T3 fibroblasts decreased histone H3K9me3 levels (Fig. 6B) as observed in aging. However, other age-associated epigenetic markers such as lower H3K27me3 and higher H4K20me1 were not observed (Fig. 6C, D, Fig. S10A, B). SUN1 WT, but not SUN1 C759A, induced age-related mitochondrial changes, including decreased membrane potential and increased mitochondrial superoxide and reactive oxygen species (ROS) (Fig. 6E-G). These results indicate that elevated SUN1 induces aging hallmarks in fibroblasts and support a role for LINC complex-dependent mechanotransduction in mediating cellular aging.

**Fig. 6.**
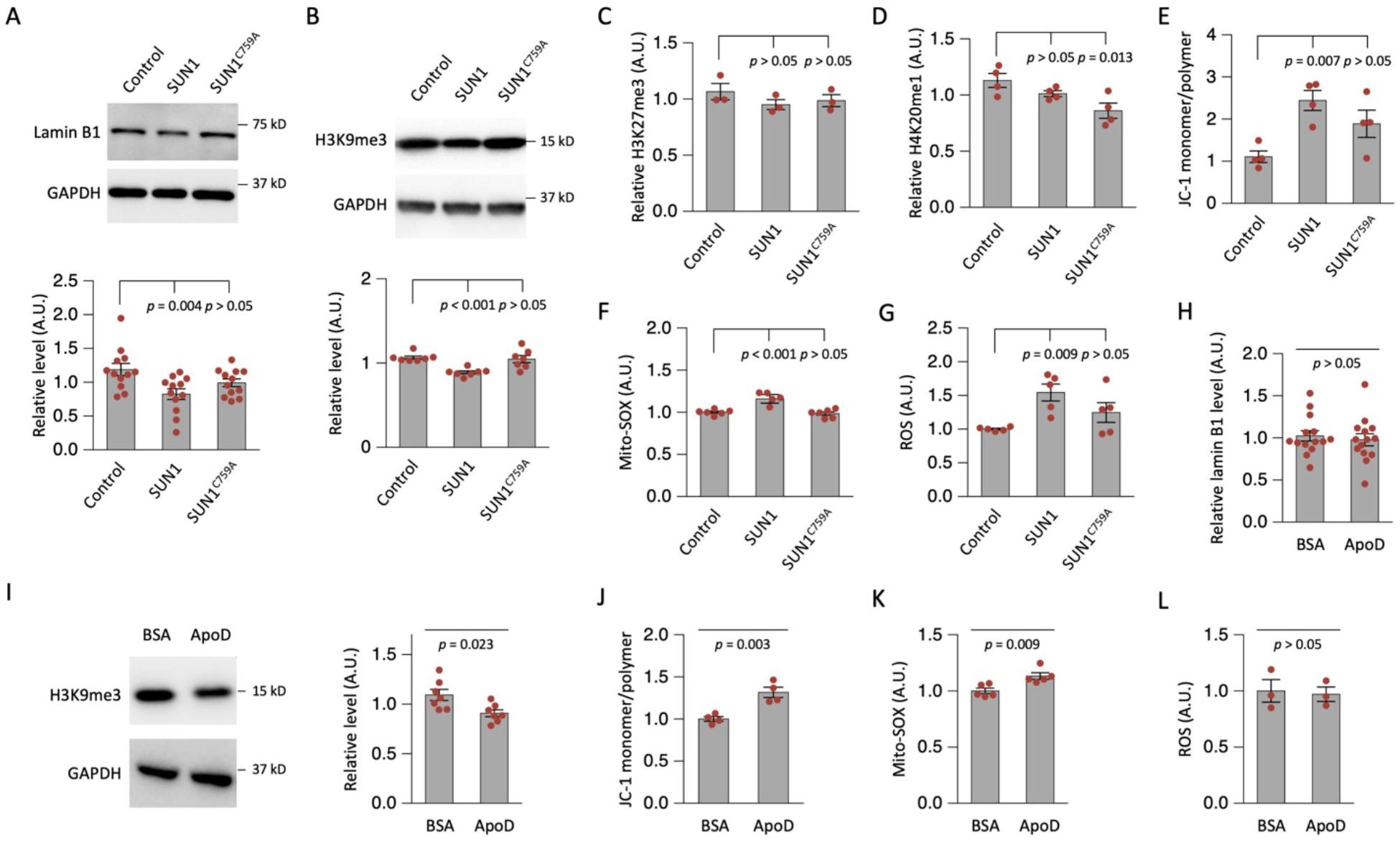
SUN1 and ApoD promote cellular aging phenotypes. (**A**) Representative Western blot and quantification of lamin B1 in NIH3T3 fibroblasts expressing myc-SUN1 or myc-SUN1^C759A^. (**B**) Representative immunoblot and quantification of H3K9me3 in cells expressing myc-SUN1 or myc-SUN1^C759A^. (**C**-**G**) Quantification of H3K27me3 (C), H4K20me1 (D), mitochondrial membrane potential (E), mitochondrial superoxide (SOX, F), and cellular ROS (G) in cells expressing the indicated myc-tagged proteins. (**H**) Lamin B1 expression after 1 day of ApoD treatment (0.1 µg/mL). (**I**) Representative immunoblot and quantification of H3K9me3 in ApoD-treated cells. (**J**-**L**) Mitochondrial membrane potential (J), mitochondrial SOX (K), and cellular ROS (L) in control and ApoD-treated cells. Values are mean ± SEM (*N* ≥ 3 experiments). *p* values were calculated with 1-way ANOVA with post hoc Dunnett’s test (A-G) or unpaired *t-*test (H-L).

We also examined the above parameters in NIH3T3 fibroblasts treated with ApoD for one day. Lamin B1 expression was unaffected by this treatment (Fig. 6H, S10C). Similar to SUN1 overexpression, ApoD treatment caused a reduction in H3K9me3 (Fig. 6I), without affecting the levels of H3K27me3 or H4K20me1 (Fig. S10D, E). ApoD treatment also caused a loss of mitochondrial membrane potential and a small increase in mitochondrial superoxide (Fig. 6J, K) but did not significantly change total ROS levels (Fig. 6L). Taken together, stable SUN1 overexpression and short-term ApoD treatment produced comparable outcomes, inducing cellular aging markers beyond defective cell polarity.

### ApoD interferes with myotube formation *in vitro* and muscle repair *in vivo*

Muscle repair capacity declines during aging^59^. We previously found that myoblasts and fibroblasts share the same LINC complex-dependent nuclear positioning pathway to generate polarity for migration and that this polarity is critical for myoblast differentiation and myotube formation^60^. Reducing the elevated expression of SUN1 in *Lmna* mouse models of premature aging improves bone and skeletal muscle function and extends lifespan^14^, suggesting a detrimental effect of overexpressed SUN1 in muscle biology. We confirmed this with cultured C2C12 myoblasts in which stable expression of SUN1, but not SUN2, inhibited the formation of multinucleated myotubes (Fig. 7A, B).

**Fig. 7.**
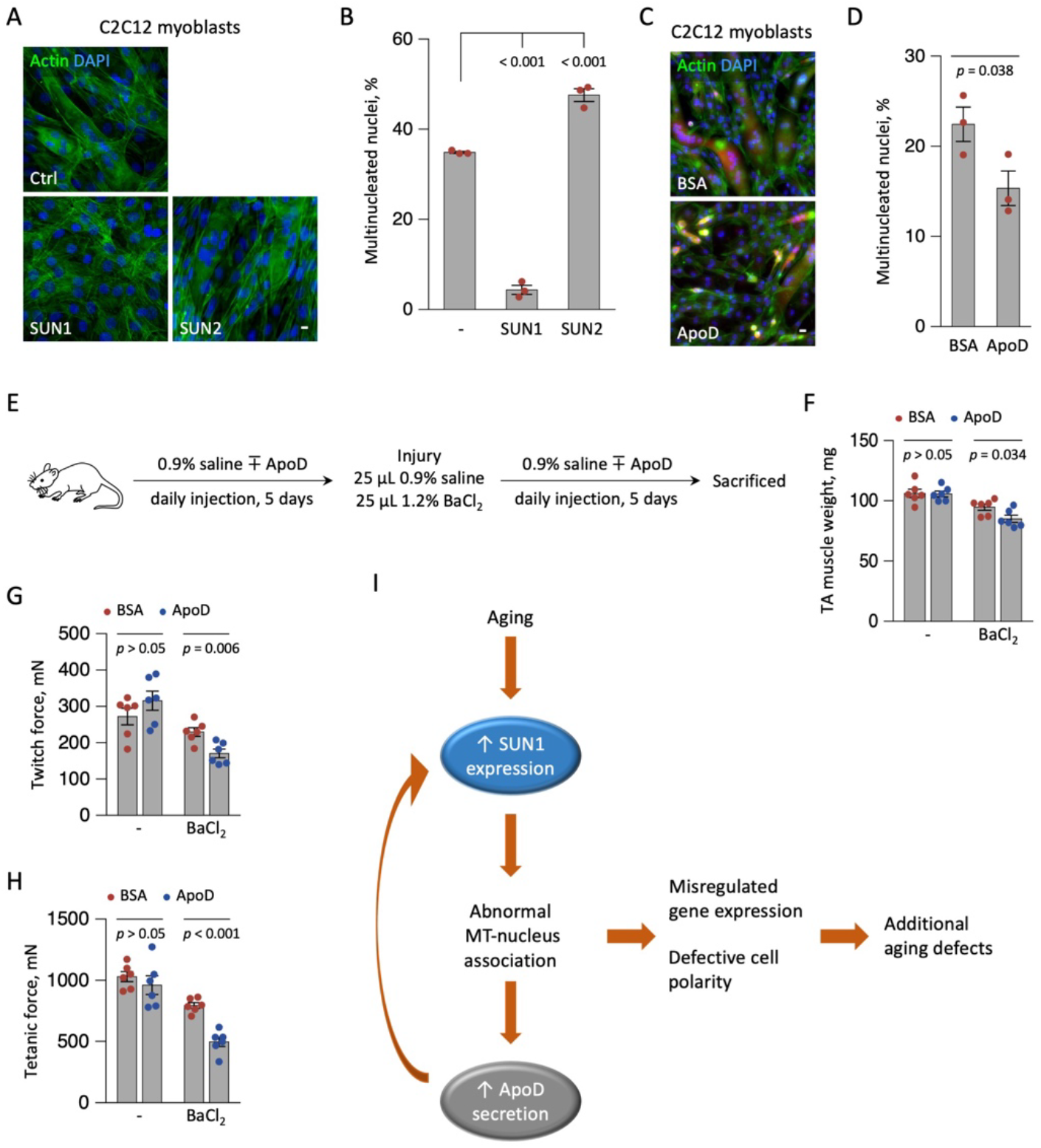
SUN1 and ApoD interfere with myotube formation. (**A, B**) Representative images (A) and quantification of fusion index (B) of C2C12 myoblasts expressing the indicated myc-tag proteins following induction of differentiation. (**C, D**) Representative images (C) and fusion index (D) of C2C12 myoblasts treated with ApoD following induction of differentiation. (**E**) Diagram of the *in vivo* muscle repair assay. (**F**-**H**) Weight (F), twitch force (G), and tetanic force (H) in TA muscles of control and ApoD-treated mice. (**I**) Model of SUN1-MT-mediated regulation of cell polarity and cellular aging: Aging-associated upregulation of SUN1 promotes aberrant interactions between the nucleus and MTs, leading to altered nuclear mechanotransduction. Disrupted nuclear mechanics in turn impair cell polarity and gene regulation, creating a feedback loop that further elevates SUN1 expression and drives progressive aging-related defects. Bars are 10 µm. Values are mean ± SEM (B, D: *N* = 3 experiments; F-H: *N* = 6 animals). *p* values were calculated with unpaired *t-*test.

We next studied the response to ApoD treatment in C2C12 myoblasts. We confirmed that SUN1 expression in myoblasts was upregulated by ApoD (Fig. S11A), mirroring its effect on SUN1 expression in fibroblasts (Fig. 3E). Myoblast differentiation into myotubes *in vitro* was significantly reduced by ApoD (Fig. 7C, D). To investigate the effect of ApoD on muscle repair *in vivo*, we utilized a barium chloride (BaCl_2_)-induced muscle injury mouse model^61^ (Fig. 7E). Mice were divided into four groups, with half of the animals receiving daily intravenous injection of ApoD for 10 d and the other half injected with an equal amount of BSA as a control. A BaCl_2_ solution was injected into each leg of half of the animals in each group on d 5 to induce muscle damage. Tibialis anterior (TA) muscle weight and function were assessed on d 10^62^. ApoD did not affect body weight (Fig. S11B) but significantly reduced the weight of the TA (Fig. 7F), the ratio of TA weight to body weight (Fig. S11C) and the TA muscle cross-sectional area upon muscle injury (Fig. S11D). ApoD treatment also reduced repair of muscle function after BaCl_2_-induced damage as measured by muscle twitch force (single stimulus) (Fig. 7G) and tetanic contraction force (sustained stimulus, 100 Hz) (Fig. 7H). In uninjured mice, ApoD treatment did not significantly affect body weight, TA muscle weight, cross-sectional area, or muscle function (Fig. 7G, H, Fig. S11B-D). Thus, ApoD interferes with muscle repair without affecting uninjured muscle *in vivo*.

## Discussion

Circulating factors have attracted growing interest in anti-aging research because heterochronic parabiosis experiments strongly implicate blood-borne molecules as causal regulators of aging^63^. Although exposure to young blood has been shown in some cases to rejuvenate aged tissues^63^, a study found that perfusion with saline-albumin could produce comparable benefits^64^. Other studies show that circulating pro-aging factors may be as or even more responsible than anti-aging factors in contributing to organismal aging^65^. We now show that ApoD secretion by aged fibroblasts forms a positive feedback loop with elevated SUN1 levels to enhance the connection of MTs to the nucleus, leading to impaired cell polarity and aging defects (Fig. 7I). This loop provides a mechanistic basis for our observation that cell polarity in HFs becomes defective after age 60^10^. This may provide an explanation for the recent finding that aging is not a linear process and includes a period of accelerated aging at age 60^33,34^. A critical issue is how this feedback loop becomes activated. In premature aging, our current results and those of others suggest that expression of progerin or other forms of unprocessed prelamin A can trigger it. In physiological aging, the trigger is currently unclear. Inflammation is known to result in ApoD release^55^ and perhaps persistent age-associated inflammation triggers ApoD accumulation.

One of the most unexpected conclusions from our study is that there is a direct MT-SUN1-nesprin-2 LINC complex pathway that regulates the expression of ApoD and other genes. This pathway requires a covalent linkage between SUN1 and nesprin and requires the MT functionality of nesprin-2, suggesting the possibility that forces generated by MT motors through the LINC complex may directly affect gene expression. There is precedent for MT force transmission across the nuclear envelope in meiosis, where MT-driven forces move synapsing chromosomes attached to the LINC complex through their protein complexes at their ends^66–68^. Virtually all previous studies suggested that actin mechanotransduction to the nucleus is responsible for regulating gene expression^26,69^. Yet, for all the nesprin-2-regulated genes whose expression was rescued by the expression of one of the nesprin-2 fragments (191 in total), 93 require only the MT-interacting fragment, whereas 96 require both actin and MT functionalities of nesprin-2. As far as we are aware, this latter finding represents the first instance in which both cytoskeletal functionalities of a nesprin are required. Our findings suggest that SUN1-MT interactions may couple mechanical cues to chromatin reorganization, potentially accounting for the mechanically driven changes in genome organization previously reported in skin HFs^70,71^.

ApoD is a lipocalin family member expressed by astrocytes, fibroblasts, and other cell types. Its plasma levels increase with age and serve as biomarkers for neurological disorders and sarcopenia^48,72^. ApoD-transgenic mice exhibit robust neuroprotection and enhanced resistance to cellular stress; however, they develop metabolic abnormalities with age^49^. By approximately 9 months of age, these mice display hepatic steatosis and subsequently develop glucose intolerance associated with insulin resistance^73,74^. Despite these metabolic phenotypes, no difference in baseline lifespan has been reported^75^. In dermal fibroblasts, ApoD expression rises during aging and contributes to diminished wound healing in the skin^76,77^. The circulating nature of ApoD allows it to affect distant tissues where it alters cellular functions through SUN1 and MTs. With its well-established role in intercellular lipid shuttling, ApoD may also regulate lipid metabolism, such as arachidonic acid (AA), and thereby modulate inflammatory signaling. Curiously, AA is itself produced by a nuclear mechano-responsive pathway in which cPLA2 is activated by expansion of the nuclear membrane^78^. It will be important to explore potential cross-talk between the cPLA2-AA and the LINC complex-MT nuclear mechano-pathways. We anticipate that the LINC-microtubule network will emerge as a central hub for mechanotransduction.

## Acknowledgments

We thank the Mass Spectrometry Core Facility of Columbia University and the Biological Imaging and Stem Cell Core of University of Macau for technical support; and Dr. Brain Burke for providing the mouse SUN1 antibodies.

## Author contributions

Conceptualization: WC, GGG, MC

Methodology: WC, MC, WY, PCW, RLB, GGG

Investigation: WC, MC, WY, YM, YL, PCW, RLB

Formal analysis: WC, MC, WY, YM, YL, RLB

Visualization: WC, MC, YM, PCW, GGG

Funding acquisition: WC, GGG, XW, RLB, JDL

Supervision: WC, GGG, XW, JDL

Writing – WC, MC, GGG

Writing – review & editing: WC, MC, GGG, XW, JDL

## Data and materials availability

All next-generation sequencing data associated with this study were deposited in the NCBI Gene Expression Omnibus (accession GSE337762).

## References

1. López-Otín, C., Blasco, M. A., Partridge, L., Serrano, M. & Kroemer, G. Hallmarks of aging: An expanding universe. Cell 186, 243–278 (2023).

2. Serebryannyy, L. & Misteli, T. Protein sequestration at the nuclear periphery as a potential regulatory mechanism in premature aging. J. Cell Biol. 217, 21–37 (2018).

3. Pathak, R. U., Soujanya, M. & Mishra, R. K. Deterioration of nuclear morphology and architecture: A hallmark of senescence and aging. Ageing Res. Rev. 67, 101264 (2021).

4. Dahl, K. N. et al. Distinct structural and mechanical properties of the nuclear lamina in Hutchinson-Gilford progeria syndrome. Proc. Natl. Acad. Sci. U. S. A. 103, 10271–10276 (2006).

5. Verstraeten, V. L. R. M., Ji, J. Y., Cummings, K. S., Lee, R. T. & Lammerding, J. Increased mechanosensitivity and nuclear stiffness in Hutchinson-Gilford progeria cells: Effects of farnesyltransferase inhibitors. Aging Cell 7, 383–393 (2008).

6. Revêchon, G. et al. Recurrent somatic mutation and progerin expression in early vascular aging of chronic kidney disease. Nat. Aging 5, 1046–1062 (2025).

7. Olive, M. et al. Cardiovascular pathology in Hutchinson-Gilford progeria: Correlation with the vascular pathology of aging. Arterioscler., Thromb., Vasc. Biol. 30, 2301–2309 (2010).

8. Scaffidi, P. & Misteli, T. Lamin A-dependent nuclear defects in human aging. Science 312, 1059–63 (2006).

9. Kubben, N. & Misteli, T. Shared molecular and cellular mechanisms of premature ageing and ageing-associated diseases. Nat. Rev. Mol. Cell Biol. 18, 595–609 (2017).

10. Chang, W. et al. Imbalanced nucleocytoskeletal connections create common polarity defects in progeria and physiological aging. Proc. Natl. Acad. Sci. U. S. A. 116, 3578–3583 (2019).

11. Chen, Z. J. et al. Dysregulated interactions between lamin A and SUN1 induce abnormalities in the nuclear envelope and endoplasmic reticulum in progeric laminopathies. J. Cell Sci. 127, 1792–804 (2014).

12. Mattioli, E. et al. Prelamin A-mediated recruitment of SUN1 to the nuclear envelope directs nuclear positioning in human muscle. Cell Death Differ. 18, 1305–15 (2011).

13. Chen, C. Y. et al. Accumulation of the inner nuclear envelope protein Sun1 is pathogenic in progeric and dystrophic laminopathies. Cell 149, 565–577 (2012).

14. Chai, R. J. et al. Disrupting the LINC complex by AAV mediated gene transduction prevents progression of Lamin induced cardiomyopathy. Nat. Commun. 12, 4722 (2021).

15. bin Imtiaz, M. K. et al. Declining lamin B1 expression mediates age-dependent decreases of hippocampal stem cell activity. Cell Stem Cell 28, 967–977 (2021).

16. Lio, C. et al. Farnesylated prelamin A induces fibroblast polarity defects in premature aging disorders by inhibiting nesprin-2-SUN2 LINC complex function. J. Cell Sci. jcs.264488 (2026) doi:10.1242/jcs.264488.

17. Starr, D. A. & Fridolfsson, H. N. Interactions between nuclei and the cytoskeleton are mediated by SUN-KASH nuclear-envelope bridges. Annu. Rev. Cell Dev. Biol. 26, 421–44 (2010).

18. Crisp, M. et al. Coupling of the nucleus and cytoplasm: Role of the LINC complex. J. Cell Biol. 172, 41–53 (2006).

19. Luxton, G. W. G., Gomes, E. R., Folker, E. S., Vintinner, E. & Gundersen, G. G. Linear arrays of nuclear envelope proteins harness retrograde actin flow for nuclear movement. Science 329, 956–9 (2010).

20. Zhu, R., Antoku, S. & Gundersen, G. G. Centrifugal displacement of nuclei reveals multiple LINC complex mechanisms for homeostatic nuclear positioning. Curr. Biol. 27, 3097–3110.e5 (2017).

21. Gümüşderelioğlu, S. et al. The KASH protein UNC-83 differentially regulates kinesin-1 activity to control developmental stage-specific nuclear migration. Curr. Biol. 35, 4668–4683.e6 (2025).

22. Horn, H. F. et al. The LINC complex is essential for hearing. J. Clin. Invest. 123, 740–750 (2013).

23. Andreu, I. et al. Mechanical force application to the nucleus regulates nucleocytoplasmic transport. Nat. Cell Biol. 24, 896–905 (2022).

24. Hernandez, M. et al. Mechanostimulation Promotes Nuclear and Epigenetic Changes in Oligodendrocytes. J. Neurosci. 36, 806–813 (2016).

25. Seelbinder, B. et al. Nuclear deformation guides chromatin reorganization in cardiac development and disease. Nat. Biomed. Eng. 5, 1500–1516 (2021).

26. Carley, E. et al. The LINC complex transmits integrin-dependent tension to the nuclear lamina and represses epidermal differentiation. eLife 10, e58541 (2021).

27. Alam, S. G. et al. The mammalian LINC complex regulates genome transcriptional responses to substrate rigidity. Sci. Rep. 6, 38063 (2016).

28. Wong, X.Loo, T.-H. & Stewart, C. L. LINC complex regulation of genome organization and function. Curr. Opin. Genet. Dev. 67, 130–141 (2021).

29. Shokrollahi, M. et al. DNA double-strand break–capturing nuclear envelope tubules drive DNA repair. Nat. Struct. Mol. Biol. 31, 1319–1330 (2024).

30. Lei, K. et al. Inner nuclear envelope proteins SUN1 and SUN2 play a prominent role in the DNA damage response. Curr. Biol. 22, 1609–1615 (2012).

31. Procter, D. J., Furey, C., Garza-Gongora, A. G., Kosak, S. T. & Walsh, D. Cytoplasmic control of intranuclear polarity by human cytomegalovirus. Nature 587, 109–114 (2020).

32. Buglak, D. B. et al. Nuclear SUN1 stabilizes endothelial cell junctions via microtubules to regulate blood vessel formation. eLife 12, e83652 (2023).

33. Shen, X. et al. Nonlinear dynamics of multi-omics profiles during human aging. Nat. Aging 4, 1619–1634 (2024).

34. Ding, Y. et al. Comprehensive human proteome profiles across a 50-year lifespan reveal aging trajectories and signatures. Cell 188, 5763–5784.e26 (2025).

35. Ruckh, J. M. et al. Rejuvenation of Regeneration in the Aging Central Nervous System. Cell Stem Cell 10, 96–103 (2012).

36. Villeda, S. A. et al. The ageing systemic milieu negatively regulates neurogenesis and cognitive function. Nature 477, 90–94 (2011).

37. Brack, A. S. et al. Increased Wnt Signaling During Aging Alters Muscle Stem Cell Fate and Increases Fibrosis. Science 317, 807–810 (2007).

38. Loffredo, F. S. et al. Growth Differentiation Factor 11 Is a Circulating Factor that Reverses Age-Related Cardiac Hypertrophy. Cell 153, 828–839 (2013).

39. Pálovics, R. et al. Molecular hallmarks of heterochronic parabiosis at single-cell resolution. Nature 603, 309–314 (2022).

40. Palazzo, A. F. et al. Cdc42, dynein, and dynactin regulate MTOC reorientation independent of Rho-regulated microtubule stabilization. Curr. Biol. 11, 1536–1541 (2001).

41. Gomes, E. R., Jani, S. & Gundersen, G. G. Nuclear movement regulated by Cdc42, MRCK, myosin, and actin flow establishes MTOC polarization in migrating cells. Cell 121, 451–463 (2005).

42. Magdalena, J., Millard, T. H. & Machesky, L. M. Microtubule involvement in NIH 3T3 Golgi and MTOC polarity establishment. J Cell Sci 116, 743–756 (2003).

43. Borrego-Pinto, J. et al. Samp1 is a component of TAN lines and is required for nuclear movement. J. Cell Sci. 125, 1099–1105 (2012).

44. Bun, P. et al. Mechanical Checkpoint For Persistent Cell Polarization In Adhesion-Naive Fibroblasts. Biophys. J. 107, 324–335 (2014).

45. Cain, N. E. et al. Conserved SUN-KASH Interfaces Mediate LINC Complex-Dependent Nuclear Movement and Positioning. Curr. Biol. 28, 1–12 (2018).

46. Calero-Cuenca, F. J. et al. Ctdnep1 and Eps8L2 regulate dorsal actin cables for nuclear positioning during cell migration. Curr. Biol. 31, 1521–1530.e8 (2021).

47. Wang, B., Han, J., Elisseeff, J. H. & Demaria, M. The senescence-associated secretory phenotype and its physiological and pathological implications. Nat. Rev. Mol. Cell Biol. 25, 958–978 (2024).

48. Waldner, A., Dassati, S., Redl, B., Smania, N. & Gandolfi, M. Apolipoprotein D Concentration in Human Plasma during Aging and in Parkinson’s Disease: A Cross-Sectional Study. Parkinson’s Dis. 2018, 1–7 (2018).

49. Rassart, E., Desmarais, F., Najyb, O.Bergeron, K.-F. & Mounier, C. Apolipoprotein D. Gene 756, 144874 (2020).

50. Camato, R., Marcel, Y. L., Milne, R. W., Lussier-Cacan, S. & Weech, P. K. Protein polymorphism of a human plasma apolipoprotein D antigenic epitope. J. Lipid Res. 30, 865–875 (1989).

51. Li, Y. et al. Elevated SUN1 promotes migratory cell polarity defects through mechanically coupling microtubules to the nuclear lamina. Commun. Biol. 9, 9 (2026).

52. Shen, Q. et al. NAT10, a nucleolar protein, localizes to the midbody and regulates cytokinesis and acetylation of microtubules. Exp. Cell Res. 315, 1653–1667 (2009).

53. Larrieu, D., Britton, S., Demir, M., Rodriguez, R. & Jackson, S. P. Chemical inhibition of NAT10 corrects defects of laminopathic cells. Science 344, 527–32 (2014).

54. Do Carmo, S., Séguin, D., Milne, R. & Rassart, E. Modulation of Apolipoprotein D and Apolipoprotein E mRNA Expression by Growth Arrest and Identification of Key Elements in the Promoter*. J. Biol. Chem. 277, 5514–5523 (2002).

55. Do Carmo, S.Levros, L.-C. & Rassart, E. Modulation of apolipoprotein D expression and translocation under specific stress conditions. Biochim. Biophys. Acta, Mol. Cell Res. 1773, 954–969 (2007).

56. Jahed, Z., Shams, H. & Mofrad, M. R. K. A Disulfide Bond Is Required for the Transmission of Forces through SUN-KASH Complexes. Biophys. J. 109, 501–509 (2015).

57. Gonçalves, J. C., Quintremil, S., Yi, J. & Vallee, R. B. Nesprin-2 Recruitment of BicD2 to the Nuclear Envelope Controls Dynein/Kinesin-Mediated Neuronal Migration In Vivo. Curr. Biol. 30, 3116–3129.e4 (2020).

58. Kim, Y. The impact of altered lamin B1 levels on nuclear lamina structure and function in aging and human diseases. Curr. Opin. Cell Biol. 85, 102257 (2023).

59. Sousa-Victor, P., García-Prat, L., Serrano, A. L., Perdiguero, E. & Muñoz-Cánoves, P. Muscle stem cell aging: regulation and rejuvenation. Trends Endocrinol. Metab. 26, 287–296 (2015).

60. Chang, W., Antoku, S., Östlund, C., Worman, H. J. & Gundersen, G. G. Linker of nucleoskeleton and cytoskeleton (LINC) complex-mediated actin-dependent nuclear positioning orients centrosomes in migrating myoblasts. Nucleus 6, 77–88 (2015).

61. Tierney, M. T. & Sacco, A. Inducing and Evaluating Skeletal Muscle Injury by Notexin and Barium Chloride. in Skeletal Muscle Regeneration in the Mouse: Methods and Protocols (ed. Kyba, M.) 53–60 (Springer, New York, NY, 2016). doi:10.1007/978-1-4939-3810-0_5.

62. Yang, W., Gao, B., Qin, L. & Wang, X. Puerarin improves skeletal muscle strength by regulating gut microbiota in young adult rats. J. Orthop. Transl. 35, 87–98 (2022).

63. Conese, M., Carbone, A., Beccia, E. & Angiolillo, A. The Fountain of Youth: A tale of parabiosis, stem cells, and rejuvenation. Open Med. 12, 376–383 (2017).

64. Mehdipour, M. et al. Rejuvenation of three germ layers tissues by exchanging old blood plasma with saline-albumin. Aging 12, 8790–8819 (2020).

65. Yankova, T., Dubiley, T., Shytikov, D. & Pishel, I. Three-Month Heterochronic Parabiosis Has a Deleterious Effect on the Lifespan of Young Animals, Without a Positive Effect for Old Animals. Rejuvenation Res. 25, 191–199 (2022).

66. Yamamoto, A., West, R. R., McIntosh, J. R. & Hiraoka, Y. A Cytoplasmic Dynein Heavy Chain Is Required for Oscillatory Nuclear Movement of Meiotic Prophase and Efficient Meiotic Recombination in Fission Yeast. J. Cell Biol. 145, 1233–1250 (1999).

67. Phillips, C. M. & Dernburg, A. F. A Family of Zinc-Finger Proteins Is Required for Chromosome-Specific Pairing and Synapsis during Meiosis in C. elegans. Dev. Cell 11, 817–829 (2006).

68. Hiraoka, Y. & Dernburg, A. F. The SUN Rises on Meiotic Chromosome Dynamics. Dev. Cell 17, 598–605 (2009).

69. Kirby, T. J. & Lammerding, J. Emerging views of the nucleus as a cellular mechanosensor. Nat. Cell Biol. 20, 373–381 (2018).

70. Liu, H., Yuan, L., Baldi, L., Sornapudi, T. R. & Shivashankar, G. V. Compressive Forces Induce Epigenetic Activation of Aged Human Dermal Fibroblasts Through ERK Signaling Pathway. Aging Cell 24, e70035 (2025).

71. Liao, Y. et al. Chromatin accessibility regulates age-dependent nuclear mechanotransduction. Proc. Natl. Acad. Sci. U. S. A. 123, e2522217123 (2026).

72. Wu, J., He, Q. & Huang, L. Discovery of serum APOD as an early sarcopenia biomarker in older adults with low muscle mass: a cross-sectional proteomic and transcriptomic investigation. PeerJ 14, e21058 (2026).

73. Do Carmo, S., Fournier, D., Mounier, C. & Rassart, E. Human apolipoprotein D overexpression in transgenic mice induces insulin resistance and alters lipid metabolism. Am. J. Physiol. Endocrinol. Metab. 296, E802–811 (2009).

74. Desmarais, F., Bergeron, K. F., Rassart, E. & Mounier, C. Apolipoprotein D overexpression alters hepatic prostaglandin and omega fatty acid metabolism during the development of a non-inflammatory hepatic steatosis. Biochim. Biophys. Acta, Mol. Cell Biol. Lipids 1864, 522–531 (2019).

75. Ganfornina, M. D. et al. Apolipoprotein D is involved in the mechanisms regulating protection from oxidative stress. Aging Cell 7, 506–515 (2008).

76. Takaya, K., Asou, T. & Kishi, K. Identification of Apolipoprotein D as a Dermal Fibroblast Marker of Human Aging for Development of Skin Rejuvenation Therapy. Rejuvenation Res. 26, 42–50 (2023).

77. Becirovic-Agic, M. et al. Faster skin wound healing predicts survival after myocardial infarction. Am J Physiol Heart Circ Physiol 322, H537–H548 (2022).

78. Enyedi, B., Jelcic, M. & Niethammer, P. The Cell Nucleus Serves as a Mechanotransducer of Tissue Damage-Induced Inflammation. Cell 165, 1160–1170 (2016).

