## Supplement figures and tables for "A SUN1-ApoD feedback loop promotes cellular aging via microtubule-nuclear mechanotransduction"

### Materials and Methods

#### *Reagents*

LPA was purchased from Avanti Polar Lipids, and fluorescent dye-conjugated phalloidin from Thermo Fisher Scientific. Recombinant ApoD expressed in mammalian cells was obtained from Novoprotein (cat# C556) and ProSpec-Tany (cat# cyt-547). Cycloheximide was obtained from MCE (cat# 66819). Unless otherwise noted, all other chemicals were from Sigma-Aldrich.

Rat monoclonal anti-tyrosinated α-tubulin antibody (clone YL1/2) was from the European Collection of Animal Cell Cultures. Anti-lamin A/C (sc-20681) and anti-ApoD (sc-166612) rabbit polyclonal antibodies and anti-SUN1 (sc-293292), anti-GFP (sc-9996), and anti-HP1α (sc-130446) mouse monoclonal antibodies were from Santa Cruz Biotechnology. Mouse anti-SUN1 (X12.11) monoclonal antibody was generously provided by Dr. Brian Burke (A*STAR, Singapore). Anti-histone H3 (60932SF), anti-phospho-histone H2A.X (9718S), ant-H3K27me3 (9733S), and anti-H3K9me3 (13969S) rabbit monoclonal antibodies and anti-H4K20me1 (9724S) rabbit polyclonal antibody were from Cell Signaling Technology. Anti-Rad51 (ab133534) rabbit monoclonal antibody, anti-ApoD (ab187513) rabbit polyclonal antibody, and anti-myosin skeletal heavy chain (ab51263) mouse monoclonal antibody were from abcam. Horseradish peroxidase-conjugated secondary antibodies were purchased from GE Healthcare, and dye-conjugated secondary antibodies were from Jackson ImmunoResearch.

DCFDA and JC-1 assay kits were obtained from Beyotime (S0033S, C2003S). MitoSOX Red mitochondrial superoxide indicator was from Invitrogen (M36008). pMSCV-puro-myc-SUN1, pMSCV-puro-myc-SUN1C759A, pSuper-retro-puro-NC, and pSuper-retro-puro-SUN1 plasmids were described previously[1,2].

#### *Cloning*

pLKO2-hygro H1 vector was derived from TRC2-pLKO-puro plasmid by replacing puromycin selectable marker gene and human U6 promotor with hygromycin selectable marker gene and human H1 promoter, respectively. shControl (sense oligo: 5’-CAACAAGATGAAGAGCACCAA-3') and shNesprin-2 (sense oligo: 5’-GCACGTAAATGACCTATAT-3’) were inserted in to pLKO2-hygro H1 using AgeI/EcoRI restriction enzymes. pInducer100-puro tet-on EGFP-C4 vector was derived from pInducer20 (Addgene #44012) by 1) inserting EGFP sequence downstream of the tetracycline-inducible minimal CMV promoter and 2) replacing the UbC promoter-neomycin resistance selectable marker sequence with human PGK promoter-puromycin resistance selectable marker sequence. The vector was digested with NotI restriction enzyme. Nesprin-2 fragments were inserted into the cut plasmid to create pInducer100-puro tet-on EGFP-C4 miniN2G, pInducer100-puro tet-on EGFP-C4 mNesp2-SR51-56-KASH, and pInducer100 puro tet-on EGFP-C4 mNesp2 JA'-CORE. All constructs were verified by DNA sequencing.

#### *Cell culture and drug treatment*

Serum-free medium (SFM) was defined as Dulbecco’s modified Eagle medium (DMEM; Gibco) supplemented with 1% penicillin-streptomycin (Thermo Fisher Scientific). NIH3T3 cells were maintained in SFM containing 10% bovine calf serum (Gemini). Dermal HFs were cultured in SFM supplemented with 15% fetal bovine serum (FBS; Gemini). HEK293T, RPE, and C2C12 cells were maintained in SFM containing 10% FBS. All cells were cultured at 37°C in a humidified incubator with 5% $CO_2$. For serum starvation, cells were washed twice with phosphate-buffered saline (PBS) and once with SFM, then incubated in SFM for 1-4 days.

For ApoD treatments, 0.005-0.25 µg/mL ApoD was added to the culture medium. Unless otherwise specified, ApoD was applied 24 hours before LPA stimulation. HPI-4 was used at 10 µM for 1 hour. Remodelin was used at 50 µM for 1 hour.

##### *Generation of stable cell lines*

Stable cell lines were generated using retroviral transduction. pMSCV or pSUPER expression plasmids, together with packaging plasmids encoding Gag-Pol and VSV-G, were transfected into HEK293T cells using calcium phosphate precipitation. After 6 hours, the transfection mixture was replaced with fresh medium, and virus-containing supernatants were collected at 24, 28, and 36 hours post-transfection, aliquoted and stored at -80 °C. For infection, thawed viral supernatants were added to target cells in the presence of 4 µg/mL polybrene for 24 hours. Cells were then selected with 1 µg/mL puromycin to establish stable populations.

Lentivirus was similarly produced in 293T cells and NIH 3T3 cells ($0.5 \times 10^6$ cells/12-well) were infected with 0.5 mL virus (in 0.6 mL total volume) with 4 µg/mL polybrene. Cells were selected using puromycin (1.5 µg/mL) and hygromycin B (300 µg/mL) for two passages. Protein expression via pInducer100 constructs was induced with 50 ng/ml doxycycline for 24 hours.

##### *Preparation and treatment of conditioned medium (CM)*

Serum-starved NIH3T3 fibroblasts were cultured in SFM for 24 hours, and the resulting conditioned medium (CM) was collected and filtered through 0.2 µm filters (Sartorius, #210175103). Serum-starved HFs were cultured in SFM for 48 hours before their CM was collected.

For dialysis, CM samples were placed into ultrafiltration tubes with 3 kDa or 10 kDa molecular-weight cutoffs (Sartorius, #VS0192, #VS0102) and dialyzed against 0.25 M ammonium acetate (pH 7.2) at 4 °C for 30 hours, with buffer changes every 10 hours. Ultracentrifugation was performed using a TLA-120.2 rotor (Beckman) at 100,000 ×g for 30 minutes. Heat inactivation was performed by incubating CM in a boiling water bath for 10 min. RNase A treatment (1 µg/mL; Qiagen) was carried out at 37 °C for 1 hour. For protease treatments, 2 units/mL proteinase K (New England Biolabs) or 0.005% trypsin-EDTA (Invitrogen) was added and incubated at 37 °C for 1 hour, followed by inactivation with 5 mM phenylmethylsulfonyl fluoride (PMSF; Beyotime, #ST2573) or 5 mg/mL trypsin inhibitor (Beyotime, #SG2033), respectively.

For CM treatments, recipient HFs were serum-starved for two days and washed once with SFM before CM was applied. Cell polarity was assessed two days later. For NIH3T3 fibroblasts, serum starvation was performed for 1 day, after which half of the culture medium was replaced with CM. Cell polarity was assessed 1 day later.

##### *Human plasma*

Human plasma was provided by Macao Blood Transfusion Service, Health Bureau of the Government of Macao. All donors were Asian males. (The only inclusion criteria is age ≤ 25 years for young individuals and age ≥ 65 for aged individuals. Informed consent was obtained from all participants included in the study. This study was approved by the Panel on Research Ethics of University of Macau Research Committee (BSERE22-APP008-FHS).

For treatments, NIH3T3 fibroblasts were serum-starved for 1 day and 2% v/v of plasma was added. LPA-stimulated cell polarization was assessed 1 day latter.

##### *Immunofluorescence staining and microscopy*

Cells were fixed with 4% paraformaldehyde (Electron Microscopy Sciences) for 10 min. Coverslips were then washed three times with PBS and blocked for 30 min in blocking buffer (PBS containing 0.5% Triton X-100 and 2% normal donkey serum; Jackson ImmunoResearch). Primary staining was performed by incubating coverslips for 1 hour at room temperature with YL1/2 antibody (1:40), mouse anti-SUN1 antibody (1:100), and/or rabbit anti-SUN1 antibody (1:200) diluted in blocking buffer. After three 5-min washes with PBS, cells were incubated with dye-conjugated phalloidin (Thermo Fisher Scientific), 4',6-diamidino-2-phenylindole (DAPI), and/or secondary antibodies (1:200) in blocking buffer.

Following staining, coverslips were washed three additional times with PBS and mounted onto glass slides using Fluoromount-G (SouthernBiotech). Images were acquired using a Nikon Ti2 microscope equipped with a 40× Plan-Apo objective (N.A. 1.0), a Zeiss LSM710 confocal microscope with a 40×/1.4 oil objective, or a Tomocube HT-2H microscope.

##### *Cell polarization assay*

Monolayers of NIH3T3 fibroblasts and HFs grown on glass coverslips were serum-starved for 2 days and 4 days, respectively, and then scratched with a pipette tip to create a wound. After recovery for 30 min, cells were stimulated with 10 µM LPA to induce polarization. Cells were fixed with 4% paraformaldehyde 2 hours after stimulation and stained as described above.

Images were acquired using a Nikon Ti2 microscope equipped with a 40× Plan-Apo objective (N.A. 1.0). Nuclear positions were quantified using CellPlot[3], with at least 30 nuclei analyzed per condition. The extent of centrosome orientation was determined from analyses of at least 60 cells per condition.

##### *Western Blot*

Cells were lysed in RIPA buffer, and protein concentrations were quantified using the BCA Protein Assay Kit (Thermo Fisher Scientific) following the manufacturer's instructions. Equal amounts of protein were resolved by SDS-PAGE and transferred onto nitrocellulose membranes. Membranes were blocked in TBS-T containing 5% non-fat milk and then incubated with primary antibodies overnight at 4 °C, followed by incubation with secondary antibodies at room temperature for 1 hour. Protein signals were developed using ECL substrate (Bio-Rad) and imaged on a ChemiDoc MP system (Bio-Rad). Whole scans of all Western blots and protein gels were included (Fig. S12-14).

For quantification, band intensities were measured using Fiji (https://fiji.sc/), with target protein signals normalized to corresponding loading controls. To enable comparison across experiments, all normalized values were further scaled to the mean value within each experiment.

##### *qRT-PCR*

Total RNA was extracted using TRIzol reagent (Ambion). Briefly, 200 µl of chloroform was added to 1 ml of TRIzol, followed by a 3-min incubation at room temperature. The mixture was centrifuged at 12,000 rpm for 15 min at 4 °C to separate the aqueous phase. An equal volume of isopropanol was added to precipitate RNA at 4 °C, and samples were centrifuged again at 12,000 rpm for 15 min. The resulting RNA pellet was washed three times with 70% ethanol and dissolved in RNase-free water.

Reverse transcription was performed using the Bio-Rad cDNA synthesis kit (#1708891) with 500 ng of RNA as input. Priming was carried out at 25 °C for 5 min, reverse transcription at 46 °C for 20 min, and enzyme inactivation at 95 °C for 1 min. qPCR analysis was conducted with the Bio-Rad SYBR Green system (#1725120) on an Applied Biosystems 7500 instrument.

The primers used were: SUN1 Forward-1 (CAAGCTGAAAGCAGGCAGCA), SUN1 Reverse-1 (TGAGCTGCACAGTCTCCTGG), SUN1 Forward-2 (AAGCTGAAAGCAGGCAGCAG), SUN1 Reverse-2: (TGAGCTGCACAGTCTCCTGG), ApoD Forward-1 (TCACCACAGCCAAAGGACAAA), ApoD Reverse-1 (CGTTCTCCATCAGCGAGTAGT), ApoD Forward-2 (AAGTCCAGTTCTTCCCGTTGATG), ApoD Reverse-2 (CCACATGGAAGAGCCAGAAGAA), GAPDH Forward (TCAACGGGAAGCCCATCA), and GAPDH Reverse (CTCGTGGTTCACACCCATCA).

##### *Mass spectrometry*

CM from HFs was collected two days after serum-starvation. Collected CM was clarified and subsequently dialyzed against triethylammonium bicarbonate buffer using a low MWCO membrane to remove small molecules. After three buffer exchange, samples were concentrated through polyethylene glycol-based volume reduction. A total of 50-150 µg of protein from each sample was submitted to the Mass Spectrometry Core Facility at Columbia University for Tandem Mass Tag (TMT)-based quantitative proteomic analysis.

##### *ROS, MitoSOX, and mitochondrial membrane potential detection*

Reactive oxygen species (ROS) levels were measured using 2’,7’-dichlorofluorescein diacetate (DCFH-DA, Beyotime) following the manufacturer’s instructions. Briefly, cells were trypsinized and incubated with diluted DCFH-DA at 37 °C for 20 min, then washed three times with PBS. Fluorescence was detected using the FITC channel on a flow cytometer.

Mitochondrial superoxide was assessed using the MitoSOX Red indicator. After digestion, cells were incubated with 1 μM MitoSOX Red working solution at 37 °C for 30 min, washed three times with PBS, and analyzed by flow cytometry using the APC channel.

Mitochondrial membrane potential was evaluated using the JC-1 assay kit according to the manufacturer's instructions. Cells were incubated with JC-1 working solution at 37 °C for 20 min and washed three times. Following trypsinization, JC-1 monomer and polymer fluorescence signals were measured by flow cytometry, and the monomer:polymer ratio was calculated using FITC/PE channels in FlowJo (FlowJo LLC).

##### *Wound healing assay*

Cells were serum-starved for 24 hours and then subjected to a scrape-wound using 200 μl pipette tips. After wounding, cells were washed three times to remove debris, and medium containing 1% calf serum was added to promote migration. ApoD was added at the time of wounding. Images were captured at 0, 12, and 24 hours using an EVOS M7000 microscope and quantified using Fiji software.

##### *C2C12 differentiation*

C2C12 cells were cultured to confluence, after which the medium was replaced with SFM containing 2% horse serum (Gibco), with or without ApoD. Fresh differentiation medium was replenished every two days. After 8 days, cells were fixed and stained with anti-myosin heavy chain antibody (used at 1:200), DAPI, and phalloidin. The number of nuclei within multinucleated myotubes and the total number of nuclei were quantified, and the fusion index was calculated as the percentage of nuclei residing in multinucleated cells.

##### *Animal experiments*

Twenty-four 3-month-old C57BL/6J male mice were used in the study. The mice were randomly and blindly assigned into two groups of twelve. One group received a daily intravenous injection of 0.375 μg BSA in 0.9% saline for 10 consecutive days, while the other group received an equivalent daily dose of ApoD. On day five, 100 μL of 1.2% $BaCl_2$ was intravenously administered to half of the animals within each group to induce muscle injury; the remaining animals received a saline control injection.

At the experimental endpoint, the tibialis anterior (TA) muscles from both hindlimbs were collected. The right TA muscle was subjected to an *in situ* contraction assay prior to muscle collection, after which both muscles were weighed. All procedures were approved by the Animal Experimentation Ethics Committee of the Shenzhen Institute of Advanced Technology, Chinese Academy of Sciences (Reference No. SIAT-IRB-170305-YGS-WXL-A0314).

##### *In situ contraction test*

Specific twitch and tetanic forces were measured as previously described[4]. *In situ* muscle contractions were assessed using a muscle functional testing system (1305A, Aurora Scientific Inc., Newmarket, Canada). After isolation, each muscle was allowed to stabilize for 15 min before being activated twice with tetanic contractions (100 mA, 300 ms, 120 Hz), with a 5 min interval between stimuli. Optimal muscle length was determined when the force responses from the two stimulations were equivalent. Twitch force was recorded using a 100 mA, 0.2 ms pulse, and tetanic force was measured using 100 mA stimulation at 150 Hz for 500 ms.

##### *RNA extraction, sequencing, and analysis*

Total RNA was extracted with Direct-zolTM RNA MiniPrep Plus Kit (Zymo Research Corp). Library preparation was performed by the University of Florida (UF) Interdisciplinary Center for Biotechnology Research (ICBR) using an Illumina RNAseq library prep kit with ribodepletion (Illumina, San Diego, CA). Sequencing was performed on a NovaSeq6000 S4 with a 2 x 150 bp read length kit (Illumina, San Diego, CA). Short reads were filtered and trimmed using Trimmomatic[5]. QC on the original and trimmed reads was performed using FastQC. The reads were aligned to the transcriptome using STAR[6]. Transcript abundance was quantified using RSEM (v 1.3.1)[7]. Differential expression analysis was performed in ExpressAnalyst[8] using DESeq2[9], with an FDR-corrected P-value threshold of 0.05 and log2 fold change of ±0.58496. Rescue analysis was done in R Studio using custom scripts. Syne2 encoding nesprin-2 was removed from the differentially expressed genes results before subsequent analyses. Enriched Gene Ontology (GO) analysis of differentially expressed genes was performed using the Enrichr web server. We evaluated the Biological Process and terms with an adjusted $p < 0.05$ were considered statistically significant. Normalized ATAC-seq bigWig files were visualized in Integrative Genomics Viewer (IGV, v2.19.4) to compare chromatin accessibility at the Apod locus.

##### *ATAC Sequencing and Data Analysis*

Nuclei were isolated fresh from 200,000 cells and ATAC-seq libraries were prepared from aliquots of 50,000 nuclei using an ATAC-seq library preparation kit from Active Motif (Carlsbad, CA). Pooled libraries were sequenced on a Novaseq 6000 S4 2x150 flow cell (ICBR Next-Gen Sequencing Core, U Florida) at a depth of 100 million reads per sample. Reads were trimmed using Trimmomatic (1), and QC on the original and trimmed reads was performed using FastQC and MultiQC[10]. The reads were aligned to the mouse genome (GRCm39) using Bowtie2[11]. ATAC peak calling was performed using MACS[12]. Differential peak analysis was performed using DASA. Peaks were annotated using TxDb.Mmusculus.UCSC.mm39.knownGene. Custom scripts were used to produce differential peak size graphs.

##### *Statistical analysis*

All experiments were performed at least three times, or in the case of young and aged HFs, twice using two independent cell strains. Independent measurements are depicted in all histograms as individual points.  Data visualization was carried out using Microsoft Excel and DotPlot (https://github.com/SciImage/DotPlot). Statistical analyses were performed with Microsoft Excel or GraphPad Prism. For comparisons between two groups, unpaired two-tailed Student's *t*-tests were used, except in Fig. S7C, where a paired two-tailed *t*-test was applied. For comparisons involving more than two groups, one-way ANOVA followed by Tukey's or Dunnett's post-hoc tests was used.

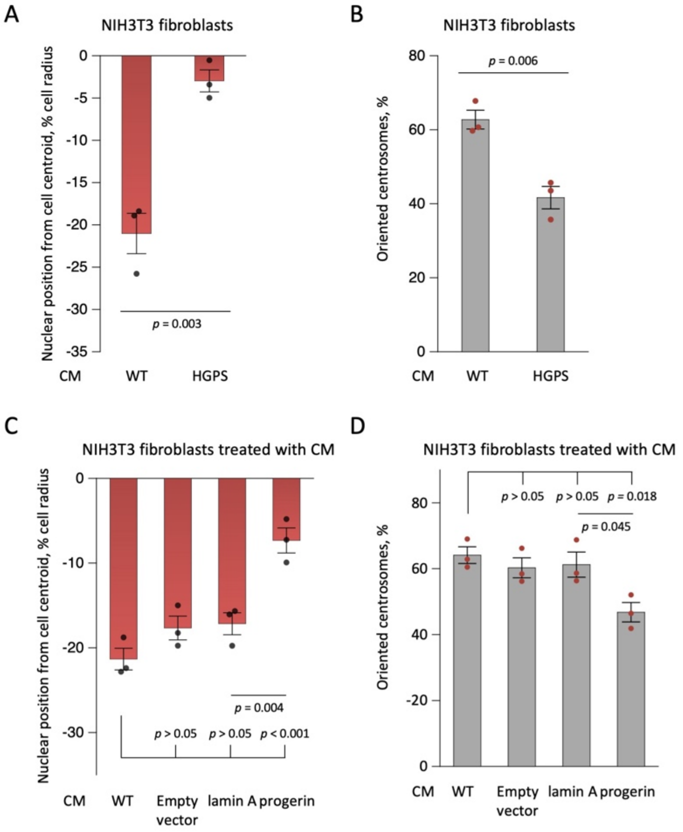


**Fig. S1. An inhibitor of cell polarity is released by cellular models of premature aging.**

(**A**, **B**) Nuclear positioning (A) and centrosome orientation (B) in LPA-stimulated NIH3T3 fibroblasts treated with CMs from HFs from control and HGPS patients. (**C**, **D**) Nuclear position (C) and centrosome orientation (D) in LPA-stimulated NIH3T3 fibroblasts treated with CMs from NIH3T3 cells expressing no plasmid (WT), empty pMSCV vector (MSCV), pMSCV-lamin A, or pMSCV-progerin. Values are mean ± SEM ($N = 3$ experiments, $n \geq 90$ cells). $p$ values were calculated with unpaired two-tailed $t$-test (A, B) or 1-way ANOVA with Tukey's test (C, D).

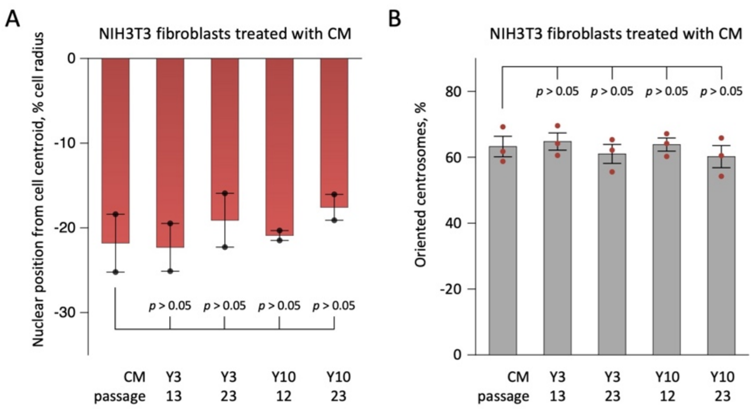


**Fig. S2. The polarity inhibitor secreted by aged HFs is not related to the senescence-associated secretory phenotype.**

Nuclear positioning (**A**) and centrosome orientation (**B**) in LPA-stimulated NIH3T3 fibroblasts treated with CMs from HFs from two young individuals. At passage 23, HFs became senescent[1], but their CM did not inhibit cell polarity in NIH3T3 fibroblasts. Values are mean ± SEM (A: $N = 2$ experiments, $n \geq 60$ cells; B: $N = 3$ experiments, $n \geq 180$ cells). $p$ values were calculated with 1-way ANOVA with Dunnett's test.

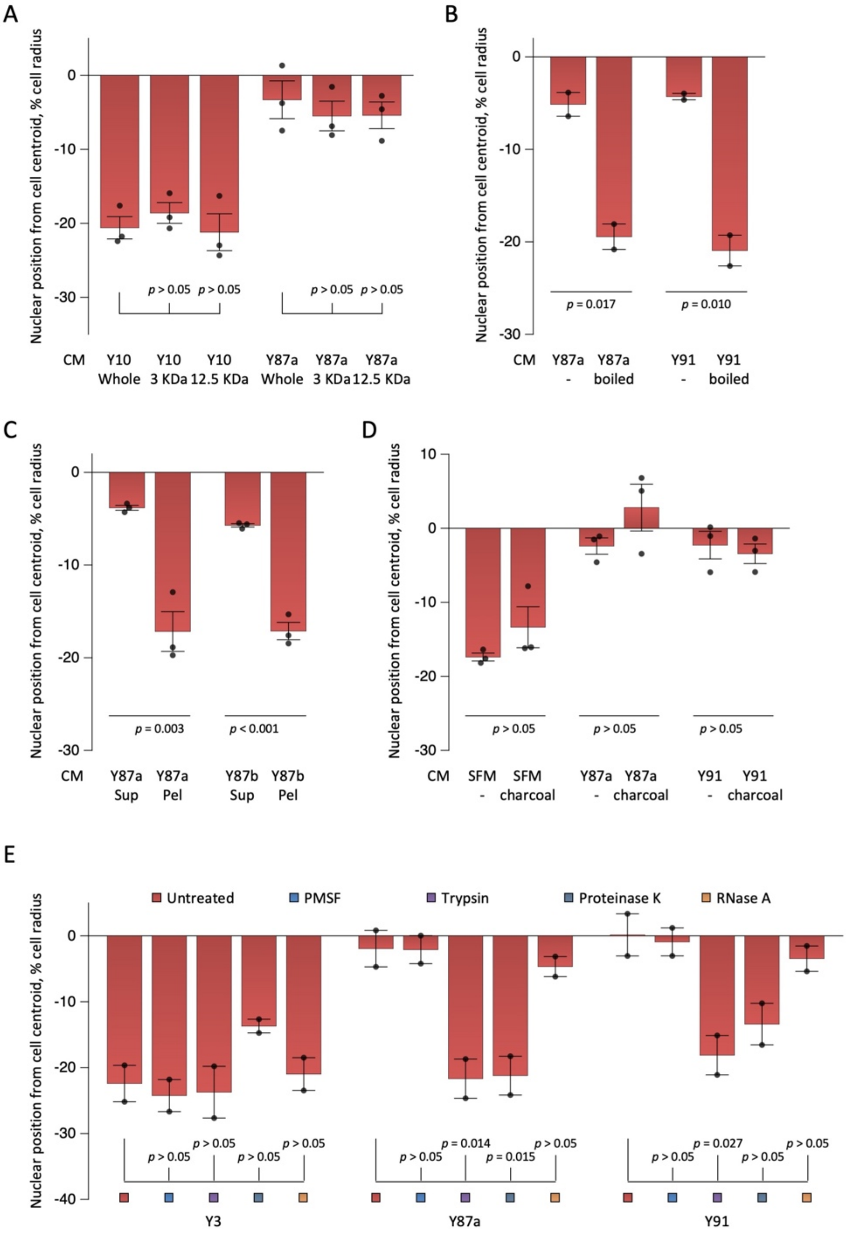


**Fig. S3. Biochemical characterization of the polarity inhibitory activity in CM from aged HFs indicated it is proteinaceous.**

Nuclear position of LPA-stimulated NIH3T3 fibroblasts treated with CM subjected to the following biochemical treatments: (**A**) Dialysis using membrane with 3 kDa or 12 kDa molecular weight cutoff; (**B**) Heat inactivation (100 ºC, 10 min); (**C**) Ultracentrifugation (100,000×*g*, 30 min) (Sup, supernatant; Pel, resuspended pellet); (**D**) Charcoal adsorption; (**E**) Enzymatic digestion using trypsin, proteinase K, and RNase A. Values are mean ± SEM (A, C, D: $N = 3$ experiments, $n \geq 90$ cells; B, E: $N = 2$ experiments, $n \geq 60$ cells). $p$ values were calculated with 1-way ANOVA with Tukey's test (A, E) or unpaired two-tailed $t$-test (B-D).

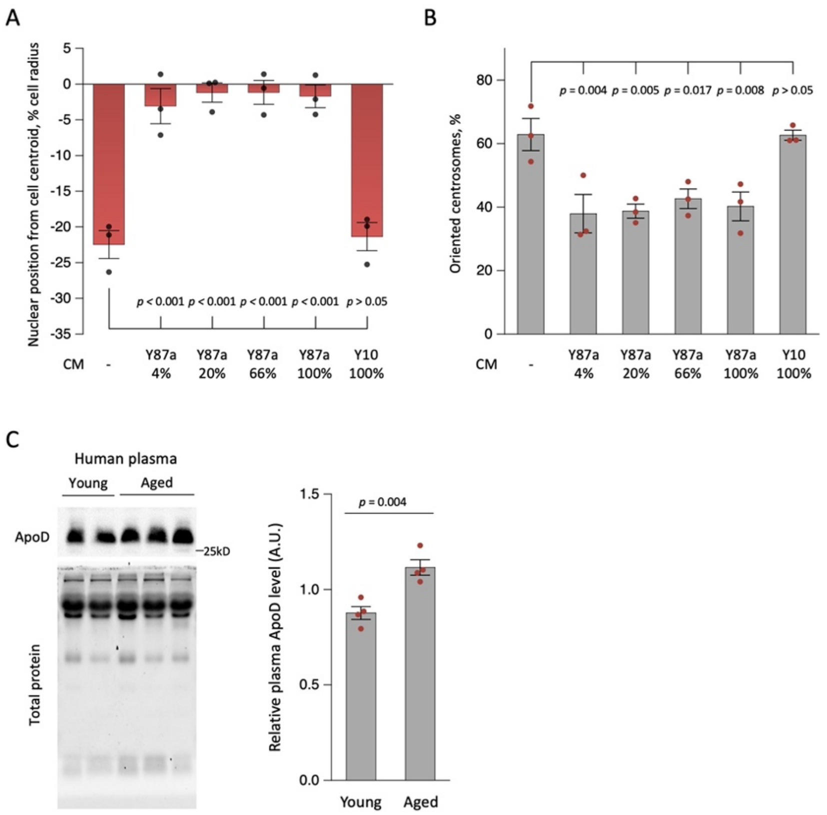


**Fig. S4. Dosage effect of CM from an aged HF.**

(**A, B**) Nuclear position (A) and centrosome orientation (B) in LPA-stimulated NIH3T3 fibroblasts treated with CM from an aged HF diluted as indicated. **(C)** Western blot (left) and quantification (right) of ApoD in plasma from young (≤ 25 yrs) and aged (≥ 65 yrs) donors. Values are mean ± SEM (A, B: $N = 3$ experiments, $n \geq 90$ cells; C: $N = 4$ experiments; $n \geq 2$ subjects per experiment). $p$ values were calculated with 1-way ANOVA with Dunnett's test (A, B) or unpaired t-test (C).

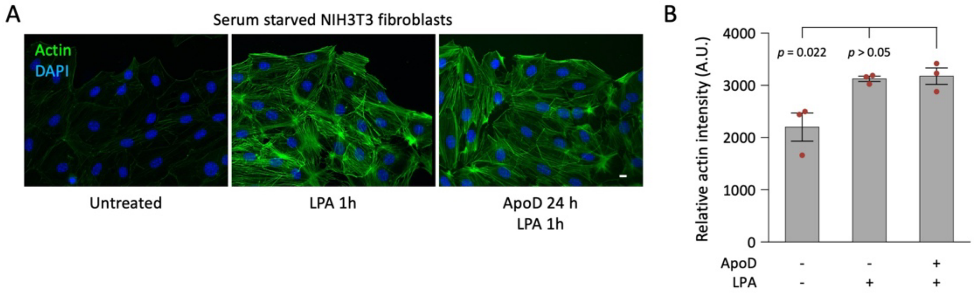


**Fig. S5. ApoD does not inhibit LPA induction of actin stress fibers.**

(**A**) Representative images of serum-starved NIH3T3 fibroblasts treated as indicated and stained for F-actin (phalloidin) and nuclei (DAPI). (**B**) Quantification of F-actin fluorescence in NIH3T3 fibroblasts treated as in (A) and stained by phalloidin. Note that LPA induced robust F-actin in serum-starved cells in both the presence and absence of ApoD (0.1 µg/mL). Values are mean ± SEM ($N = 3$ experiments). $p$ values were calculated with 1-way ANOVA with Tukey's test.

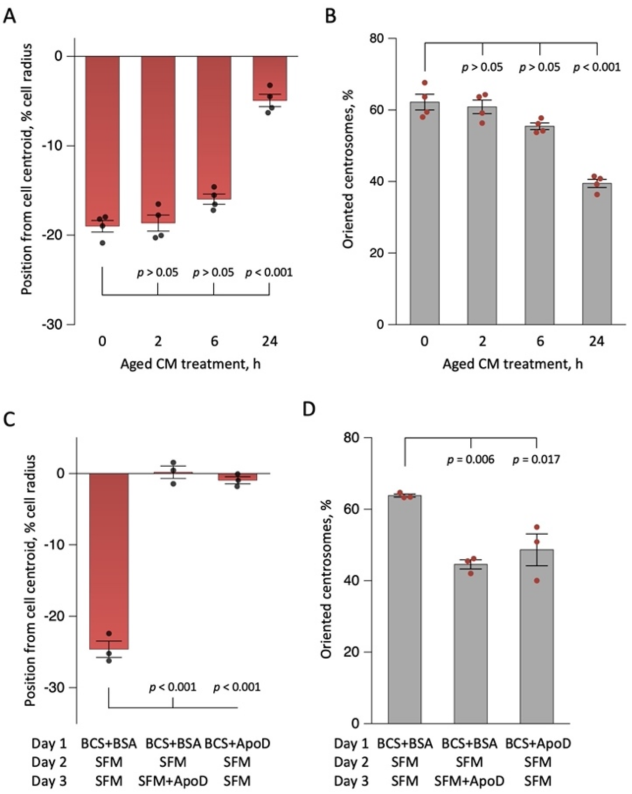


**Fig. S6. Inhibition of cell polarity by ApoD requires prolonged treatment.**

(**A**, **B**) Nuclear position (A) and centrosome orientation (B) in LPA-stimulated NIH3T3 fibroblasts treated with CM from aged HFs for the indicated durations. (**C, D**) Nuclear position (C) and centrosome orientation (D) in LPA-stimulated NIH3T3 fibroblasts treated as indicated. Centrosome orientation was inhibited when ApoD was added in the presence of serum and removed during the two days of serum starvation. Values are mean ± SEM ($N \geq 3$ experiments, $n \geq 90$ cells). $p$ values were calculated with 1-way ANOVA with Dunnett's test (A, B) or Tukey's test (C, D).

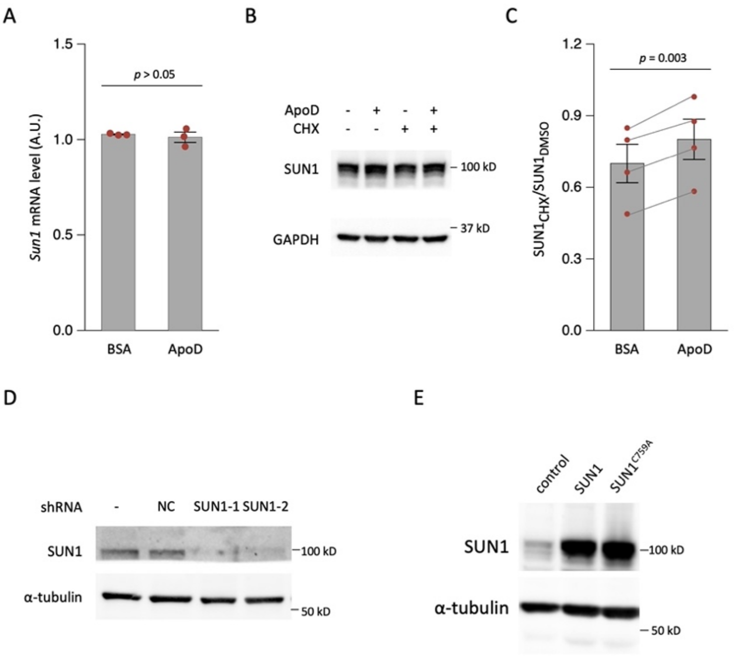


**Fig. S7. ApoD stabilizes SUN1 protein level.**

(**A**) Expression of SUN1 mRNA in NIH3T3 cells treated with 0.1 µg/mL BSA or ApoD for 1 day. (**B**) Representative immunoblot analysis of SUN1 in NIH3T3 fibroblasts treated with or without ApoD (0.1 µg/mL) and then treated with 500 µM cycloheximide (CHX) for 6 h. (**C**) Quantification of CHX experiments as in B. (**D**) Immunoblot showing knockdown of SUN1 in NIH3T3 fibroblasts by two SUN1 shRNAs; NC is noncoding shRNA control. (**E**) Immunolot showing the expression of myc-SUN and myc-SUN1$^{C759A}$ in NIH3T3 fibroblasts. Values are mean ± SEM (A: $N = 3$ experiments; C: $N = 4$ experiments). $p$ values were calculated with unpaired two-tailed $t$-test (A), paired two-tailed $t$-test (C).

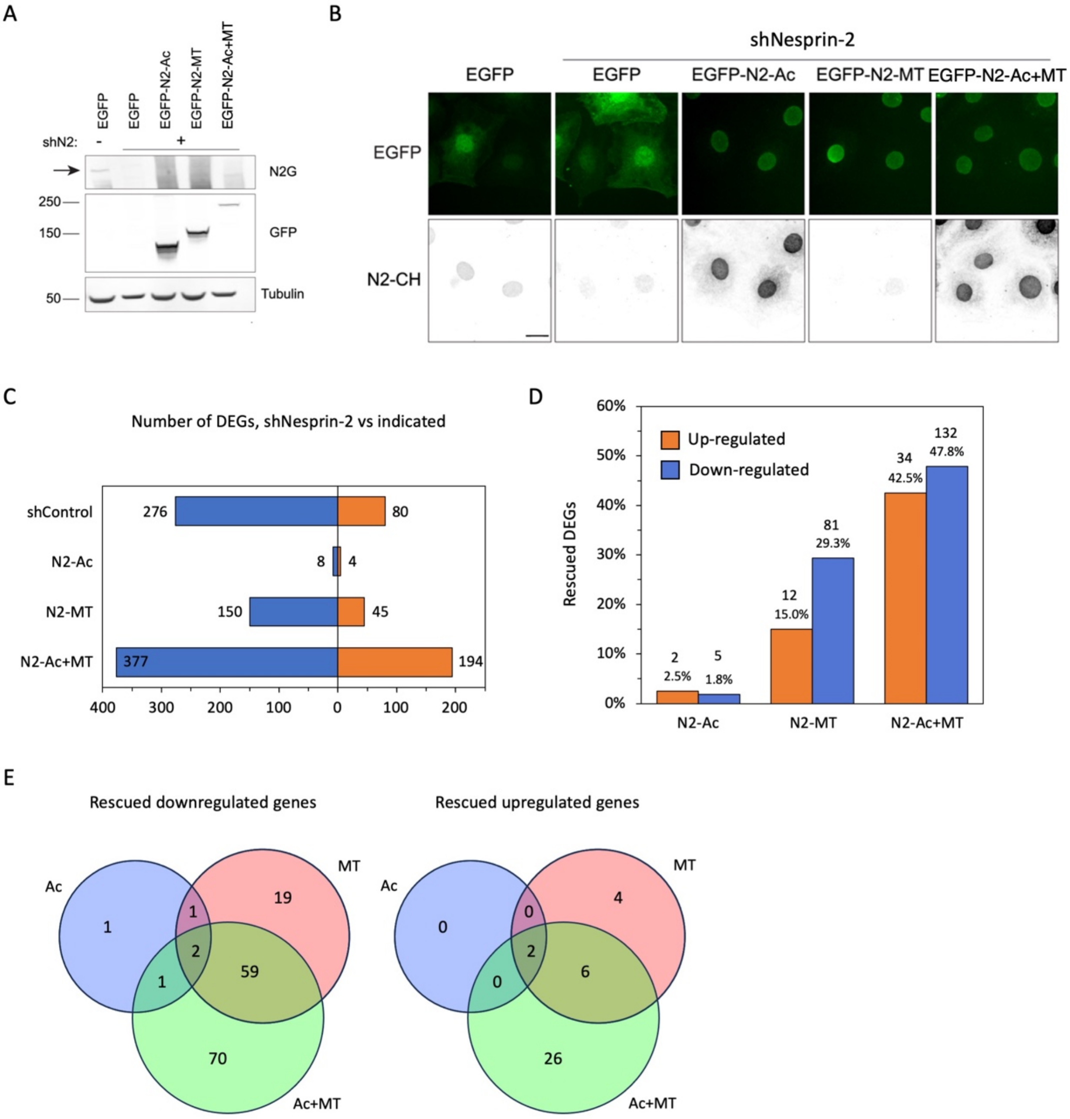


**Fig. S8. RNA-Seq analysis of gene expression in nesprin-2 depleted cells re-expressing EGFP- nesprin-2 fragments.**

(**A**) Western blots showing the depletion of nesprin-2 by an shRNA (shN2) and the expression of the GFP-tagged nesprin-2 fragments. (**B**) Representative images of GFP and immunostaining of nesprin-2 using an antibody recognizing the actin-binding calponin homology (N2-CH) domains. Bar, 20 µm. (**C**) Numbers of upregulated (orange) and downregulated (blue) differently regulated genes (DEGs) in NIH3T3 fibroblasts knocked down for nesprin-2 (shNesprin-2) vs. cells expressing the control shRNA (shControl), or shNesprin-2 and the indicated GFP-tagged nesprin-2 fragments. (**D**) Number (upper labels) and percentage (lower labels) of DEGs in C that are rescued by reexpression of the indicated GFP-N2 fragments. (**E**) Diagram showing the number of genes downregulated (left) and upregulated (right) by shN2 that are rescued by the indicated nesprin-2 fragments.

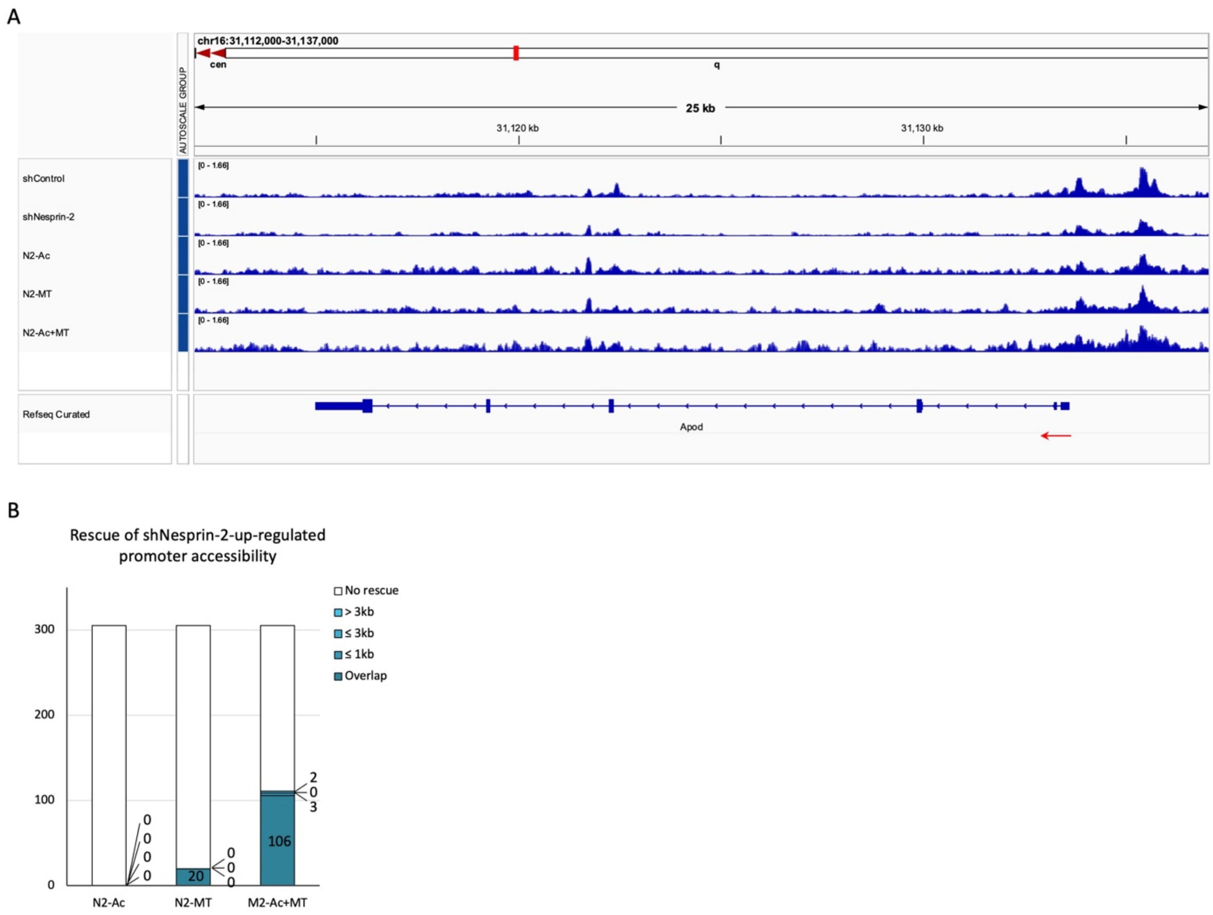


**Fig. S9. ATAC-Seq analysis of gene expression in nesprin-2 depleted cells re-expressing EGFP- nesprin-2 fragments.**

(**A**) Related to Fig. 5E. Diagram showing the accessibility of the *Apod* gene on Chromosome 16. Red arrow, transcription start site and direction. (**B**) Number of shNesprin-2-upregulated promoter regions whose accessibility was decreased by expression of the indicated nesprin-2 fragments. The regions were classified according to the distances between the shNeprin-2-upregulated peak and the peak abolished by the indicated nesprin-2 fragment. “Overlap” indicates direct overlap between the corresponding peaks; ≤ 1 kb, ≤ 3 kb, and > 3 kb denote the distance between corresponding peaks; “No rescue” indicates regions where accessibility was not restored.

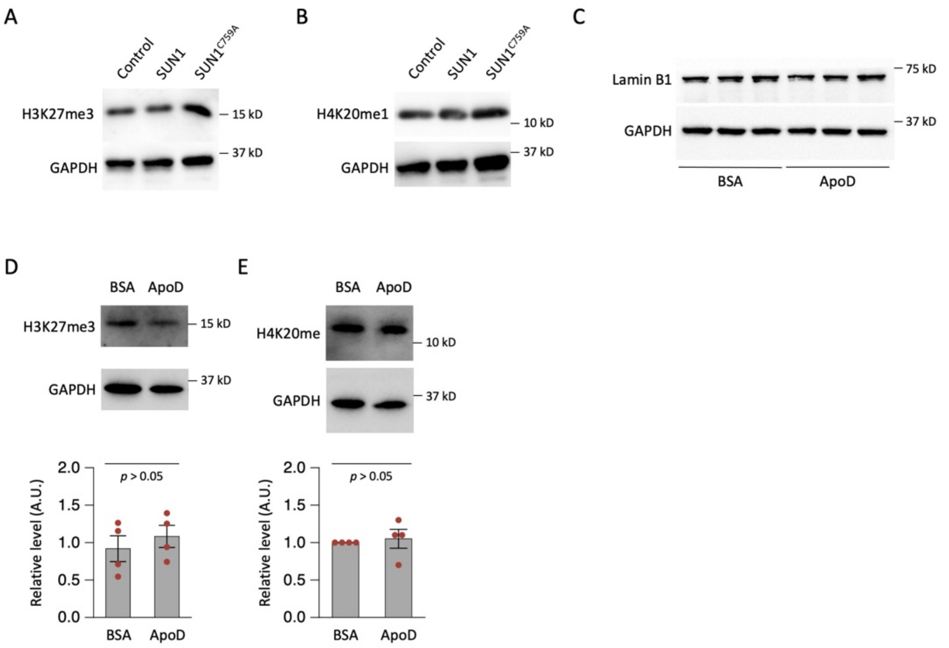


**Fig. S10. SUN1 expression and ApoD treatment induce some, but not all, features of cellular aging.**

(**A**, **B**) Representative immunoblot images of two unaffected senescence markers, H3K27me3 (A) and H4K20me1 (B), in lysates of NIH3T3 fibroblasts expressing the indicated myc-tagged proteins. (**C**) Related to Fig. 6H. Representative immunoblot of lamin B1 in lysates of NIH3T3 fibroblasts treated with either BSA or ApoD (0.1 µg/mL, 24 hrs). (**D, E)** H3K27me3 (D) and H4K20me1 (E) in lysates of NIH3T3 fibroblasts treated with either BSA or ApoD (0.1 µg/mL, 24 hrs). Values are mean ± SEM ($N = 4$ experiments). $p$ values were calculated with unpaired two-tailed $t$-test.

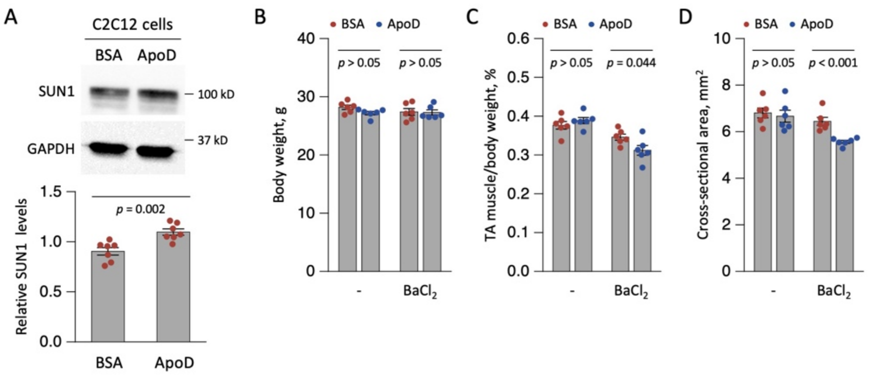


**Fig. S11. ApoD interferes with muscle repair in mice.**

(**A**) Top, representative immunoblot of SUN1 in C2C12 cells treated with ApoD (0.1 µg/mL, 24 h). Bottom, quantitation of SUN1 levels relative to GAPDH from immunoblots such as shown on the top. (**B**-**D**) Body weight (B), ratio of tibialis anterior (TA) muscle weight to body weight (C), and cross-sectional area of TA muscle (D) in mice following muscle injury and repair. Animals received daily injections of 0.375 ng of the indicated proteins for 10 days, with muscle injury induced by $BaCl_2$ injection on day 5. Tissues were collected on day 10. Values are mean ± SEM (A: $N = 7$ experiments; B-D: $N = 6$ animals). $p$ values were calculated with unpaired two-tailed $t$-test.

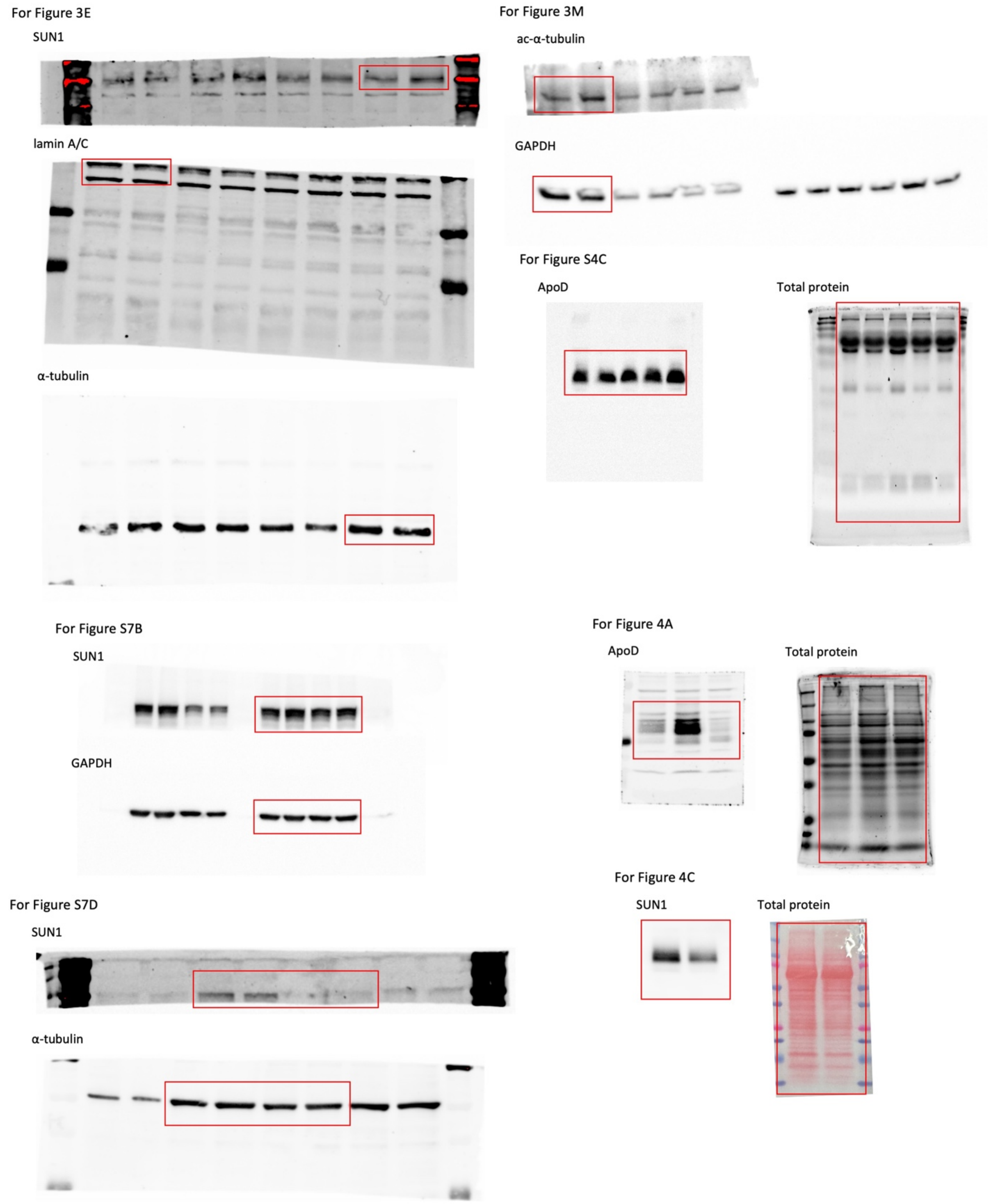


**Fig. S12. Raw images of Immunoblots.**

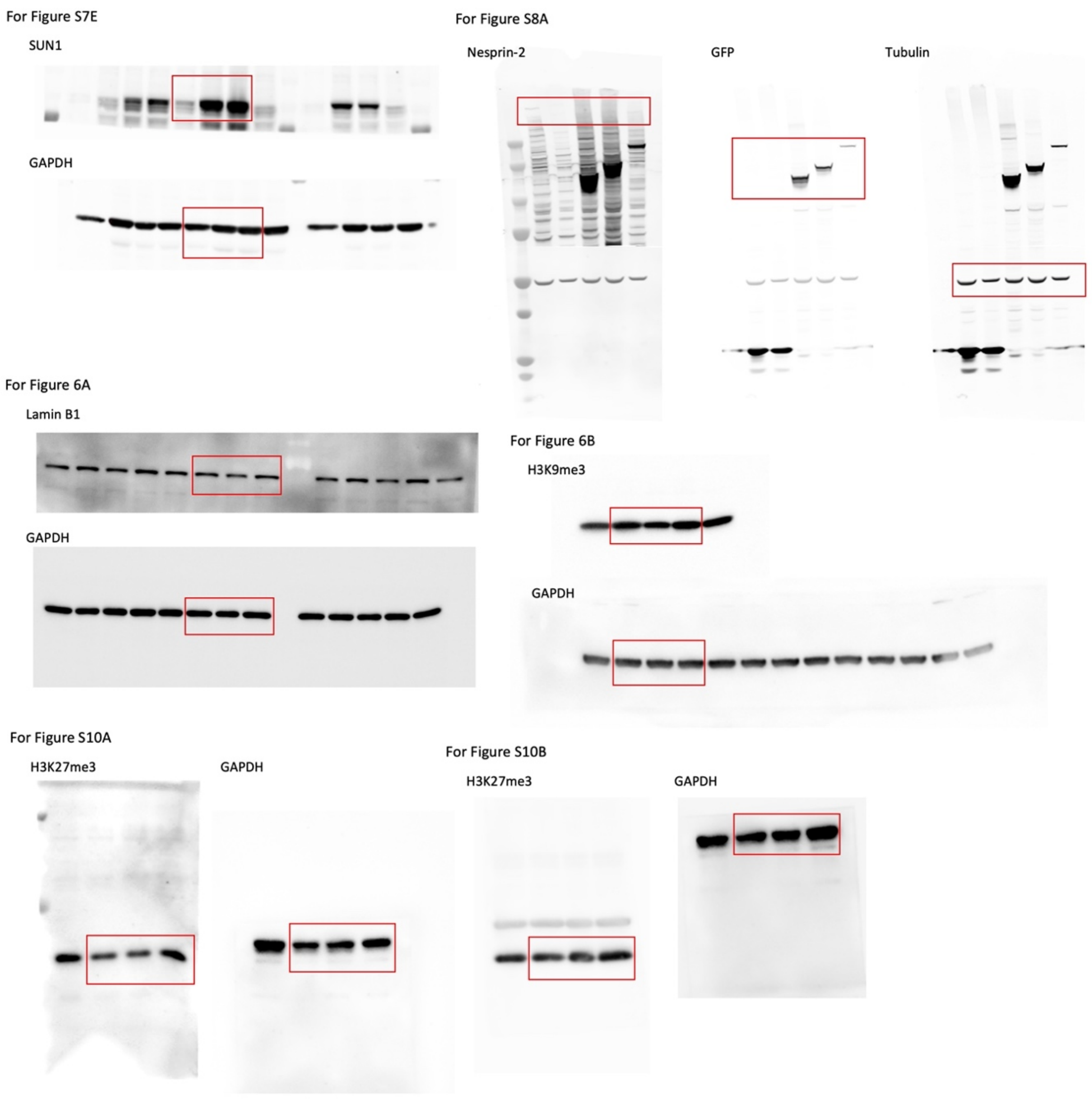


**Fig. S13. Raw images of Immunoblots.**

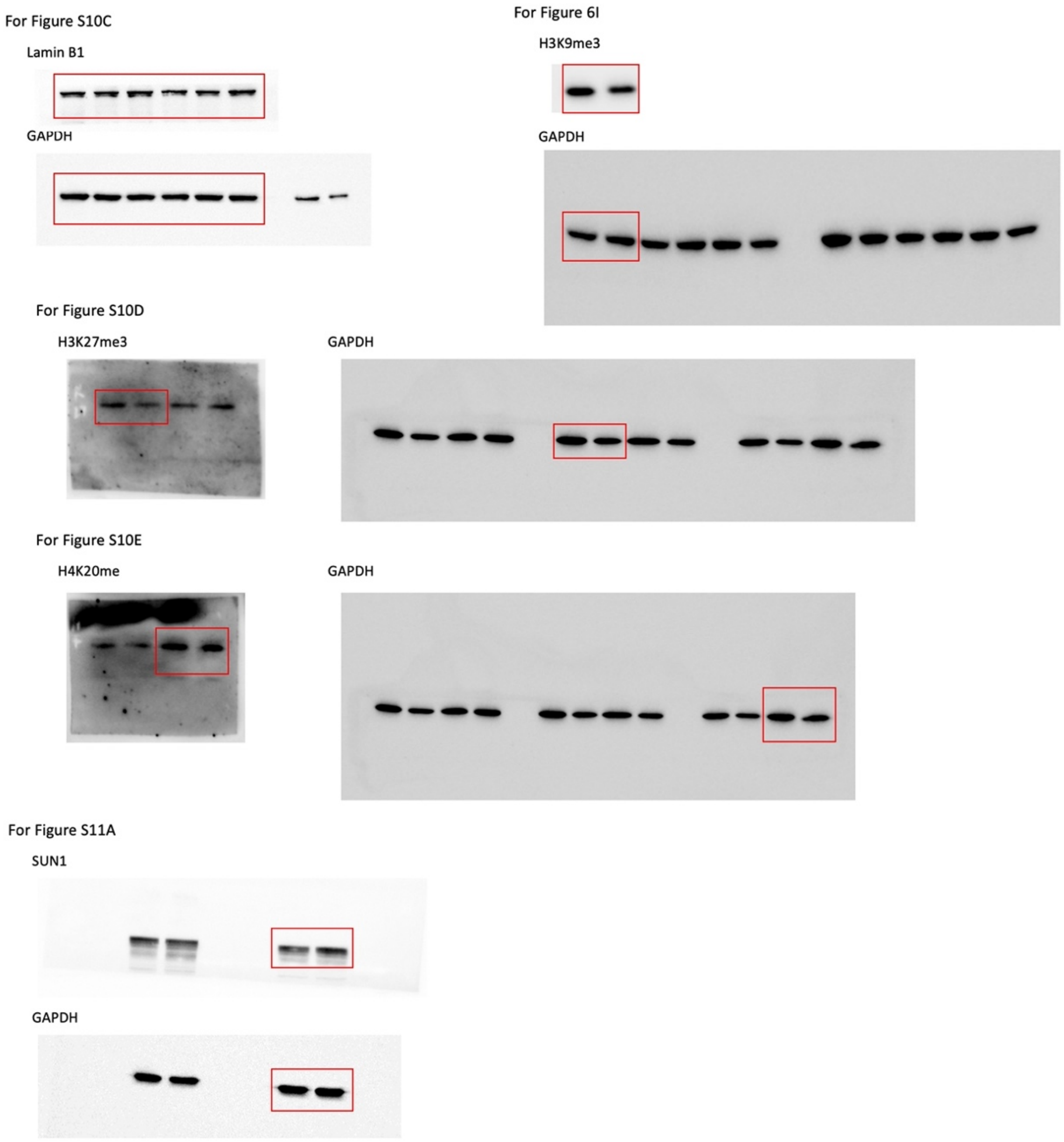


**Fig. S14. Raw images of Immunoblots.**

**Table S1. Primary human dermal fibroblasts used in this study.**

| Label | Catalog ID* | Sex | Age (year) | Note |
|---|---|---|---|---|
| | GM00316 | male | 12 | |
| | AG11498 | male | 14 | HGPS patient |
| | GM00498 | male | 3 | |
| | AG06917 | male | 3 | HGPS patient |
| | GM01652 | female | 11 | |
| | AG01972 | female | 14 | HGPS patient |
| | GM00038 | female | 9 | |
| | AG11513 | female | 8 | HGPS patient |
| **Y3** | GM05565 | male | 3 | |
| **Y5** | GM05381 | male | 5 | |
| **Y17** | AG06234 | male | 17 | |
| **Y22** | AG11747 | male | 22 | |
| **Y24** | AG11732 | female | 24 | |
| **Y30a** | AG11242 | male | 30 | |
| **Y30b** | AG13153 | male | 30 | |
| **Y59** | AG06239 | male | 59 | |
| **Y60** | AG05419 | male | 60 | |
| **Y61** | AG11011 | male | 61 | |
| **Y65** | AG04659 | male | 65 | |
| **Y68** | AG16030 | male | 68 | |
| **Y69** | AG05095 | male | 69 | |
| **Y70** | AG11733 | female | 70 | |
| **Y84** | AG05274 | male | 84 | |
| **Y87a** | AG10884 | male | 87 | |
| **Y87b** | AG05248 | male | 87 | |
| **Y91** | AG07725 | male | 91 | |

*Source: Coriell Cell Repositories.

**Table S2. Enriched Gene Ontology (GO) Biological Process terms identified using Enrichr (GO Biological Process 2026), N2-MT-rescued genes.**

| Term | Adjusted P-value |
|---|---|
| Defense Response to Virus (GO:0051607) | 7.94E-06 |
| Antiviral Innate Immune Response (GO:0140374) | 2.08E-04 |
| Regulation of Viral Genome Replication (GO:0045069) | 0.0017 |
| Regulation of Type I Interferon-Mediated Signaling Pathway (GO:0060338) | 0.0029 |
| Response to Cytokine (GO:0034097) | 0.0035 |
| Negative Regulation of Viral Genome Replication (GO:0045071) | 0.0050 |
| Negative Regulation of Type I Interferon-Mediated Signaling Pathway (GO:0060339) | 0.0060 |
| Negative Regulation of Viral Process (GO:0048525) | 0.0139 |
| Positive Regulation of Apoptotic Cell Clearance (GO:2000427) | 0.0315 |
| Regulation of Apoptotic Cell Clearance (GO:2000425) | 0.0315 |
| Positive Regulation of Interferon-Beta Production (GO:0032728) | 0.0364 |
| Interleukin-27-mediated Signaling Pathway (GO:0070106) | 0.0420 |
| Mitral Valve Morphogenesis (GO:0003183) | 0.0441 |
| Positive Regulation of Immune Response (GO:0050778) | 0.0441 |
| Complement Activation, GZMK Pathway (GO:0160257) | 0.0490 |

### References


1. Chang, W. *et al.* Imbalanced nucleocytoskeletal connections create common polarity defects in progeria and physiological aging. *Proc. Natl. Acad. Sci. U. S. A.* **116**, 3578–3583 (2019).
2. Li, Y. *et al.* Elevated SUN1 promotes migratory cell polarity defects through mechanically coupling microtubules to the nuclear lamina. *Commun. Biol.* **9**, 9 (2026).
3. Chang, W., Antoku, S. & Gundersen, G. G. Wound-healing assays to study mechanisms of nuclear movement in fibroblasts and myoblasts. in *Methods in Molecular Biology* (eds Shackleton, S., Collas, P. & Schirmer, E. C.) vol. 1411 255–267 (2016).
4. Yang, W., Gao, B., Qin, L. & Wang, X. Puerarin improves skeletal muscle strength by regulating gut microbiota in young adult rats. *J. Orthop. Transl.* **35**, 87–98 (2022).
5. Bolger, A. M., Lohse, M. & Usadel, B. Trimmomatic: a flexible trimmer for Illumina sequence data. *Bioinformatics* **30**, 2114–2120 (2014).
6. Dobin, A. *et al.* STAR: ultrafast universal RNA-seq aligner. *Bioinformatics* **29**, 15–21 (2013).
7. Li, B. & Dewey, C. N. RSEM: accurate transcript quantification from RNA-Seq data with or without a reference genome. *BMC Bioinformatics* **12**, 323 (2011).
8. Liu, P. *et al.* ExpressAnalyst: A unified platform for RNA-sequencing analysis in non-model species. *Nat Commun* **14**, 2995 (2023).
9. Love, M. I., Huber, W. & Anders, S. Moderated estimation of fold change and dispersion for RNA-seq data with DESeq2. *Genome Biol* **15**, 550 (2014).
10. Ewels, P., Magnusson, M., Lundin, S. & Käller, M. MultiQC: summarize analysis results for multiple tools and samples in a single report. *Bioinformatics* **32**, 3047–3048 (2016).
11. Langmead, B. & Salzberg, S. L. Fast gapped-read alignment with Bowtie 2. *Nat Methods* **9**, 357–359 (2012).
12. Zhang, Y. *et al.* Model-based Analysis of ChIP-Seq (MACS). *Genome Biol* **9**, R137 (2008).
